# HypVW is an HOCl-Sensing Two Component System in *Escherichia coli*

**DOI:** 10.1101/2021.09.09.459708

**Authors:** Sara El Hajj, Camille Henry, Alexandra Vergnes, Laurent Loiseau, Gaël Brasseur, Romain Barré, Laurent Aussel, Benjamin Ezraty

## Abstract

Two component systems (TCS) are signalling pathways that allow bacterial cells to sense, respond and adapt to fluctuating environments. Among the classical TCS of *Escherichia coli*, YedVW has been recently showed to be involved in the regulation of *msrPQ*, encoding for the periplasmic methionine sulfoxide reductase system. In this study, we demonstrate that hypochlorous acid (HOCl) induces the expression of *msrPQ* in a YedVW dependant manner, whereas H_2_O_2_, NO and paraquat (a superoxide generator) do not. Therefore, YedV appears to be an HOCl-sensing histidine kinase. Based on this finding, we proposed to rename this system HypVW. Moreover, using a directed mutagenesis approach, we show that Met residues located in the periplasmic loop of HypV (formerly YedV) are important for its activity. Given that HOCl oxidizes preferentially Met residues, we bring evidences that HypV could be activated via the reversible oxidation of its methionine residues, thus conferring to MsrPQ a role in switching HypVW off. Based on these results, we propose that the activation of HypV by HOCl could occur through a Met redox switch. HypVW appears to be the first characterized TCS able to detect HOCl in *E. coli*. This study represents an important step in understanding the mechanisms of reactive chlorine species resistance in prokaryotes.

**IMPORTANCE:** Understanding molecularly how bacteria respond to oxidative stress is crucial to fight pathogens. HOCl is one of the most potent industrial and physiological microbiocidal oxidant. Therefore, bacteria have developed counterstrategies to survive HOCl-induced stress. Over the last decade, important insights into these bacterial protection factors have been obtained. Our work establishes HypVW as an HOCl-sensing two component system in *Escherichia coli* MG1655 which regulates the expression of the periplasmic HOCl-oxidized proteins repair system MsrPQ. Moreover we bring evidences suggesting that HOCl could activate HypV through a methionine redox switch.

## INTRODUCTION

During infections, the host innate immune system produces high levels of reactive oxygen and chlorine species (ROS, RCS), such as hydrogen peroxide (H_2_O_2_) and hypochlorous acid (HOCl), to kill invading pathogens ^1^. One major challenge for bacteria is to overcome this oxidative stress. While bacterial responses to H_2_O_2_ have been extensively studied, much less is known about responses to HOCl. In the cell, HOCl can damage biomolecules like DNA, lipids and proteins ^2,3^. Within proteins, HOCl oxidizes effectively and preferentially sulfur-containing amino acids such as cysteine and methionine^4^. The thiol group of the cysteine can be oxidized to sulfenic (-SOH), sulfinic (-SO_2_H) and sulfonic (-SO_3_H) acids or to disulphide bond (S-S)^5^. On the other hand, the thioether side chain of methionine can be oxidized to sulfoxide, thereby, converting methionine into methionine sulfoxide (Met-O) ^6^.

Bacterial cells can prevent damages caused by HOCl through various mechanisms and pathways. Many of them are general ROS-response pathways, but others detect and respond specifically to HOCl ^7^. In *Escherichia coli*, recent studies identified several regulatory factors activated by HOCl, like the transcriptional factors HypT (a LysR-type regulator) and RclR (an AraC-type activator). HypT detects and responds to HOCl through the oxidation of three methionine residues ^8^, whereas RclR is activated through the oxidation of two cysteine residues ^9^. Upon activation, HypT up-regulates genes involved in Met and Cys biosynthesis while repressing genes implicated in iron acquisition. The activation of RclR induces the expression of *rcl* genes that are important to resist HOCl treatment ^9^. Likewise, NemR (a TetR-family repressor) functions as a HOCl-responsive transcription factor in *E. coli* that controls the expression of genes involved in HOCl resistance like *nemA* and *golA* ^10^. Moreover, repairing HOCl-damages represent another way for bacteria to survive this stress. The methionine sulfoxide reductase (MSR), which repairs HOCl-oxidized proteins, is a crucial enzyme family involved in HOCl resistance^11^.

Two-component systems (TCS) are signal transduction pathways typically made of a membrane-bound histidine kinase sensor (HK) and its cytoplasmic response regulator (RR). They sense external signals and trigger bacterial adaptive responses to a wide range of environments, stress and growth conditions ^12^. Our previous work showed that in *E. coli*, the YedVW two-component system is involved in the induction of the *msrPQ* genes in response to HOCl^13^. The *msrPQ* encodes for the periplasmic methionine sulfoxide reductase (MsrPQ), composed of a periplasmic molybdopterin-containing oxidoreductase (MsrP) and a haem-containing membrane-bound protein (MsrQ). MsrP catalyzes the reduction of protein bound Met-O and therefore repair periplasmic oxidized proteins after HOCl stress ^13^.

In the *E. coli* genome, the *yedWV* operon is located upstream and in the opposite direction as the *msrPQ* genes. The intergenic region in between contains *hiuH*, a gene encoding a transthyretin-like periplasmic protein. This protein hydrolyses 5-hydroxyisourate (5-HIU), a compound formed via the oxidation of urate ^14^. The Ishihama’s and Ogasawara’s laboratories showed that the expression of *hiuH* was induced by H_2_O_2_ in a YedVW dependent manner ^15^. Moreover, they identified the Cys165 residue located in the trans-membrane domain of YedV as being essential for the sensing of H_2_O_2_ ^16^ by oxidation of this Cys residue. Finally, they proposed to rename this system HprSR for hydrogen peroxide response sensor/regulator.

In this study, we show that *hiuH, msrP* and *msrQ* genes belong to the same operon and are under the control of YedVW. Our results point out that YedV is predominantly a RCS and not a ROS sensing histidine kinase. Using a site-specific mutagenesis approach, we report that two methionine residues located in the periplasmic loop are critical for YedV activity and that Cys165 is more likely implicated in signal transduction than signal detection. This report reveals for the first time a two-component system sensing HOCl in *E. coli*. Interestingly, it also suggests that HOCl could activate YedV through a methionine redox switch. Coherently, we propose to rename this TCS HypVW.

## RESULTS

### *hiuH, msrP* and *msrQ* belong to the same operon

In the genome of *E. coli*, the *hiuH* gene is located 109 bp and 1,114 bp upstream of *msrP* and *msrQ*, respectively (Fig. 1A). The expression of *hiuH, msrP* and *msrQ* has been reported to be under the control of the two component system YedVW ^13,16^ with a YedW box being located upstream of *hiuH* (87 to 70 bp before the *hiuH* start codon). These observations suggest that *hiuH, msrP* and *msrQ* might belong to the same operon. This hypothesis was further explored by RT-PCR using converging pairs of primers located within each of the three genes. After RNA retro-transcription, PCR amplifications were observed between *hiuH* and *msrP* and between *msrP* and *msrQ*, showing a tri-cistronic organisation (Fig. 1B). As positive and negative controls, we used chromosomal DNA and total RNA, respectively, as templates for PCR amplifications with the same pairs of primers. Together, these experiments strongly suggest that *hiuH, msrP* and *msrQ* are part of the same operon.

**Figure 1-.**
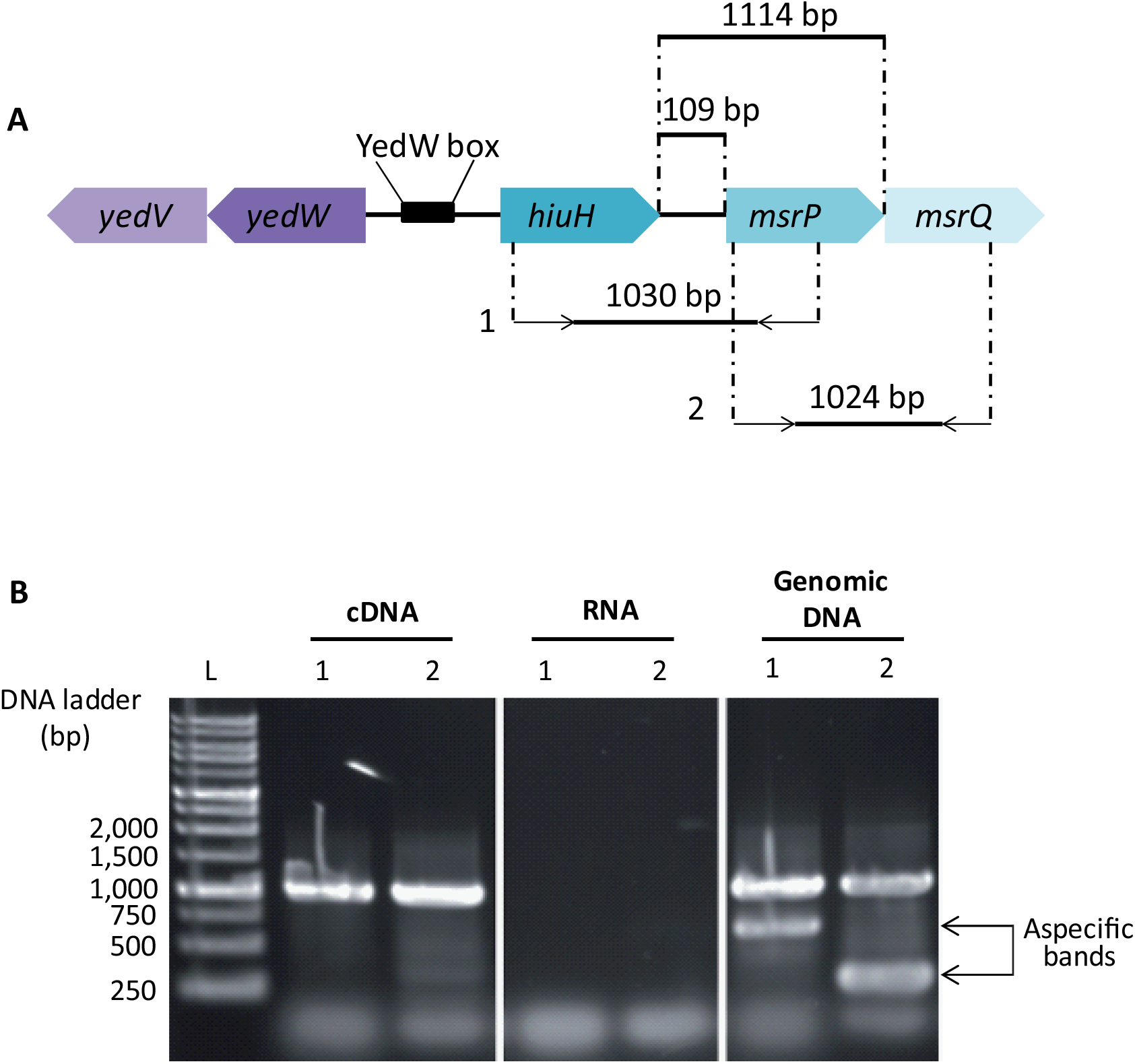
*hiuH, msrP* and *msrQ* genetic organization in *Escherichia coli* K12. **A)** Schematic representation of *yedVW* operon, *hiuH, msrP* and *msrQ* genes. In *E. coli* genome, *hiuH, msrP* and *msrQ* are adjacent. They are located in proximity to *yedV-yedW* operon. The intergenic region between *hiuH* and *yedW* contains YedW box (CATTACAAAATTGTAATG)^15^. Primers positions used for the following experiments are indicated on the scheme: the first set of primers amplify the region of 1030 pb between *hiuH* and *msrP* (1). The second set amplify the region of 1024 pb between *msrP* and *msrQ* (2) **B)** RT-PCR analysis of *hiuH, msrP* and *msrQ. E. coli* cells MG1655 (WT) were cultured in presence of 2 mM HOCl in LB medium. Total RNA was extracted and retro-transcribed to cDNA. PCR reactions were carried out on total extracted RNA, chromosomal DNA and on cDNA using the primers depicted in panel (A).

### HOCl constitute a better signal for YedVW than H_2_O_2_

To explore the physiological signals controlling the YedV-YedW regulatory system, we investigated the expression of the YedW-activated genes *hiuH, msrP, yedV and yedW* by qRT-PCR analysis and by measuring the β-galactosidase activity originating from *hiuH-lacZ* or *msrP-lacZ* translational fusions. First, we revaluated the capacity of YedVW to response to H_2_O_2_ as previously reported by Urano and collaborators^15^. We carried out qRT PCR analysis on RNAs isolated from wild type *E. coli* cells grown to mid log-phase and treated with H_2_O_2_ (Fig. 2A). The results show a 2-fold up-regulation of *yedV, yedW* and *msrP* and an approximately 7-fold up-regulation of *hiuH* in response to H_2_O_2_ treatment (6 mM as used by *Urano et al*, 2017). The induction by H_2_O_2_ was dependent on the presence of YedW since no up-regulation was observed in a Δ*yedW* mutant. However, using translational *hiuH-lacZ* or *msrP-lacZ* reporter fusions, no up-regulation was observed after H_2_O_2_ treatment in the wild-type strain (Fig. 2B). These results are in accordance with previous observations reported by Gennaris *et al*., showing that exposure of wild-type cells to H_2_O_2_ did not induce the synthesis of MsrP despite slightly inducing its transcription. However, Gennaris *et al*. reported the increase of the synthesis of MsrP after HOCl treatment^13^. Indeed, qRT PCR analysis shows that HOCl treatment (4 mM) led to a 1,500-fold and 60-fold up-regulation of *hiuH* and *msrP*, respectively, and approximately 20-fold up-regulation of *yedW* and *yedV* in the wild-type strain (Fig. 2A). Using the translational *hiuH-lacZ* and *msrP-lacZ* reporter fusions, we found a 20 and 70-fold up-regulation, respectively, after HOCl stress (Fig. 2B). Collectively, these results indicate that the induction of the *hiuH-msrPQ* operon by HOCl was dependent on the presence of a functional YedVW system. Our results also reveal that HOCl is a better signal than H_2_O_2_ for YedVW activation. Moreover, the concentration of H_2_O_2_ (6 mM) required for the activation of YedVW (at the mRNA level of *hiuH-msrPQ*) was found to be lethal (33% rate of survival) whereas the concentration of HOCl (4 mM) was nearly sub-lethal (92% survival) (Supplementary Fig. 1).

**Figure 2-.**
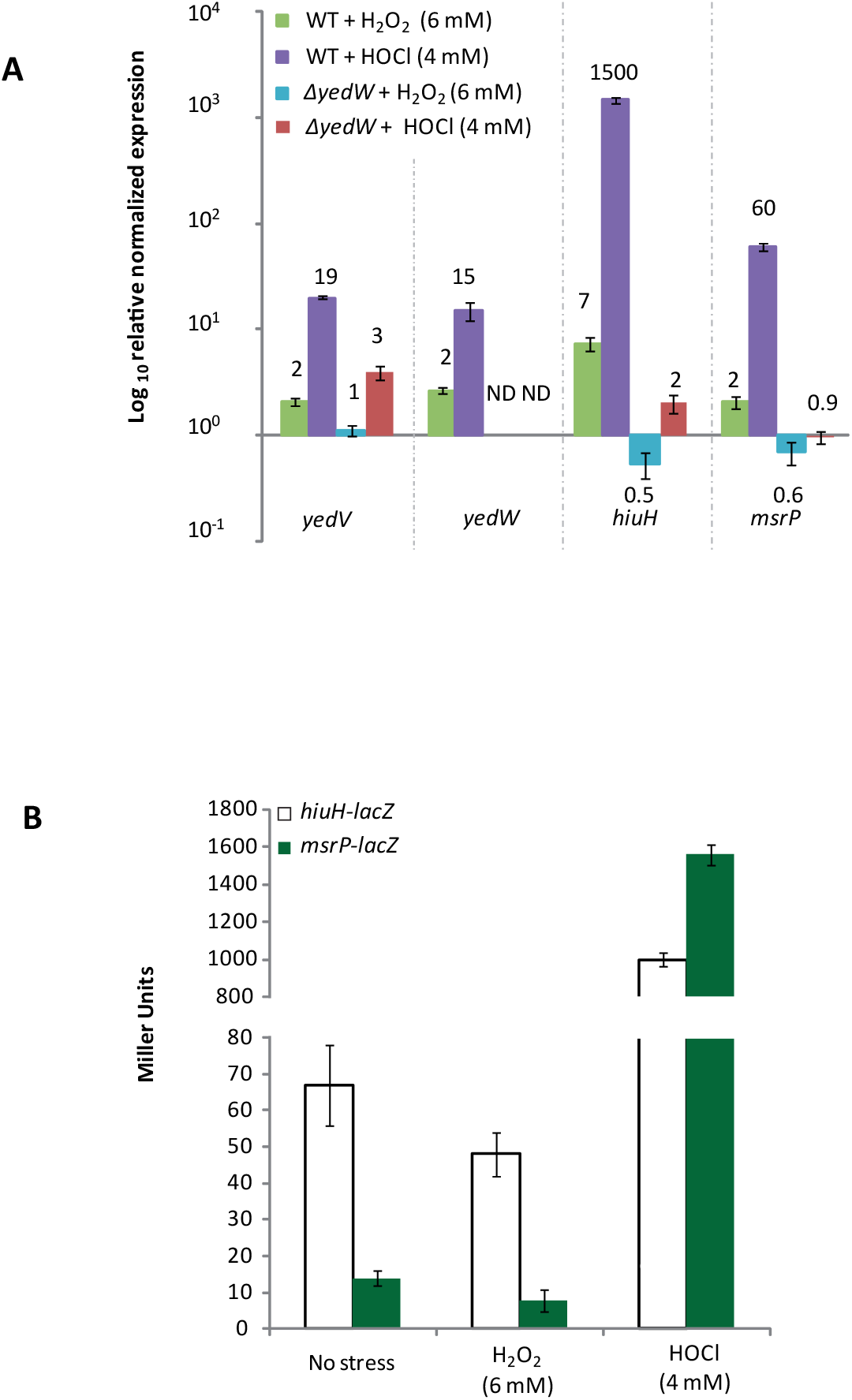
Influence of H_2_O_2_ and HOCl on the expression of the *hiuH-msrPQ* and *yedVW* operons. **A)** Fold expression level of *yedV, yedW, hiuH* and *msrP* transcripts under H_2_O_2_ and HOCl stresses in MG1655 (WT) and SH198 (*ΔyedW*) cells. Both strains are cultured in LB and subjected either to 6 mM H_2_O_2_ for 30 min, as described by Urano and his collaborators^15^, or to 4 mM HOCl for 1 hour. For each condition, quantitative real-time PCR (RT-qPCR) was performed to amplify *yedV, yedW, hiuH* and *msrP* cDNA. Values are expressed as means ± SEM of three independent experiments. ND: not detemined. **B)** Evaluation of *msrP* and *hiuH* expression under HOCl or H_2_O_2_ stress. The expression of *msrP-lacZ* and *hiuH-lacZ* fusions, in strains CH183 and CH184 respectively, were used as a read out of *hiuH* and *msrP* expression. Strains CH183 and CH184 were cultured in LB and subjected to H_2_O_2_ or HOCl stress and ß-galactosidase tests run. Results are expressed as means ± SD, n=3.

To overcome this lethality issue, different sub-lethal concentrations of H_2_O_2_ (0.5 to 3 mM) were tested with the *msrP-lacZ* expression. The H_2_O_2_ was added at OD≈0.2 and the β-galactosidase activities were measured over the course of 3 hours. For all the sub-lethal concentrations tested, no induction of the *msrP-lacZ* fusion was observed (Fig. 3A) compared to the unstressed sample, whereas in positive control experiments, the *ahpC-lacZ* fusion, which respond to H_2_O_2_ via an OxyR activation, shows a 2.5-fold induction after 30 min of exposure to 0.5 and 1 mM H_2_O_2_ (Supplementary Fig. 2A). These results challenge the relevance of the *in vivo* activation of YedVW by H_2_O_2_ and raise the question on the real signal detected by this system. Therefore, we decided to pursue the study by investigating the response of YedVW to different type of oxidants *in vivo*. We treated wild-type *E. coli* cells with various reactive oxygen, nitrogen and chlorine species at different sub-lethal concentrations. No significant expression of *msrP* was observed in response to paraquat (1,1′ -dimethyl-4,4′ -bipyridinium, a superoxide generator) or nitric oxide (NO) stress (Fig. 3B and 3C). The *soxS-lacZ* and the *hmpA-lacZ* fusions, which respond to paraquat and NO, respectively, were used as positive controls (Supplementary fig. 2B and 2C). In contrast, YedVW is activated by HOCl and *N*-Chlorotaurine (*N*-ChT), a long-lived oxidant produced by the reaction between HOCl and the amino acid taurine (Fig. 3D). Taken together, these data show that the activation of YedVW is specific to HOCl and related RCS.

**Figure 3-.**
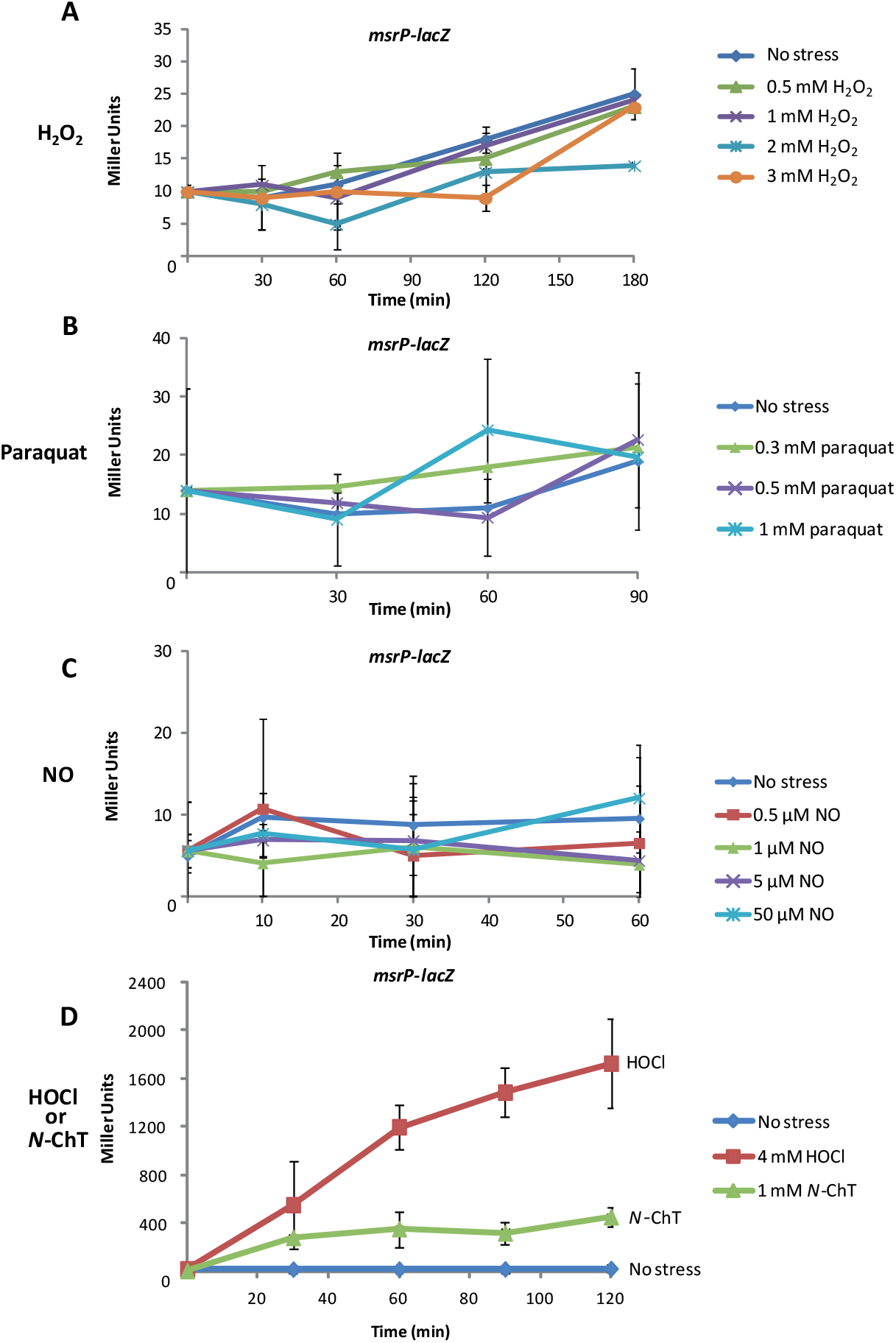
Effect of different oxidative stress on YedVW activation. The induction of *msrP-lacZ* fusion in strain CH183 is used as a read-out of YedVW activation. **A)** Effect of H_2_O_2_ on *msrP-lacZ* expression. Strain CH183 was subjected to sub-lethal concentrations of H_2_O_2_ ranging from 0.5 to 3 mM, *msrP-lacZ* expression was followed at different time points. Results are expressed as means ± SD (n=5) **B)** Effect of superoxide radical on *msrP-lacZ* expression. Paraquat was used to produce superoxide radical in the cells. Strain CH183 was subjected to sub-lethal concentration of paraquat (0.3 mM, 0.5 mM and 1 mM), *msrP-lacZ* expression was followed at different time points. Results are expressed as means ± SD (n=3) **C)** Effect of NO on *msrP-lacZ* expression. Strains CH183 was cultured anaerobically and subjected to different concentration of NO (0.5 µM, 1 µM, 5 µM and 50 µM). *msrP-lacZ* expression was followed at different time points. Results are expressed as means ± SD (n=3). **D)** Evaluation of *msrP-lacZ* expression under HOCl or *N*-ChT stress. Strain CH183 was cultured in LB and subjected to 4 mM HOCl or 1 mM *N*-ChT. ß-galactosidase assays were conducted at different time points for each condition. Results are expressed as means ± SD (n=3).

### Positions Met72 and Met153 are important for YedV activity

Our next goal was to investigate the molecular mechanism by which YedV senses HOCl. HOCl has been shown to modify proteins via the oxidation of sulfur-containing compounds, such as cysteine and methionine ^17^. Both residues have the highest degree of reactivity towards HOCl. A previous study had already investigated the implication of cysteine residues in YedV activation, and it had pointed-out the residue Cys165, located in the trans-membrane domain, as being important ^16^. As a complementary study, we decided to investigate the implication of Met residues in the function of YedV. Analysis of YedV sequence revealed the presence of 6 Met residues among which 4 (position 72, 73, 100 and 153) are located in the periplasmic loop, the first segment of YedV to be in contact with HOCl (Fig. 4A). To assess the role of each periplasmic Met residue, site directed mutagenesis with single Met-to-Ala and single Met-to-Gln substitutions were constructed.

**Figure 4-.**
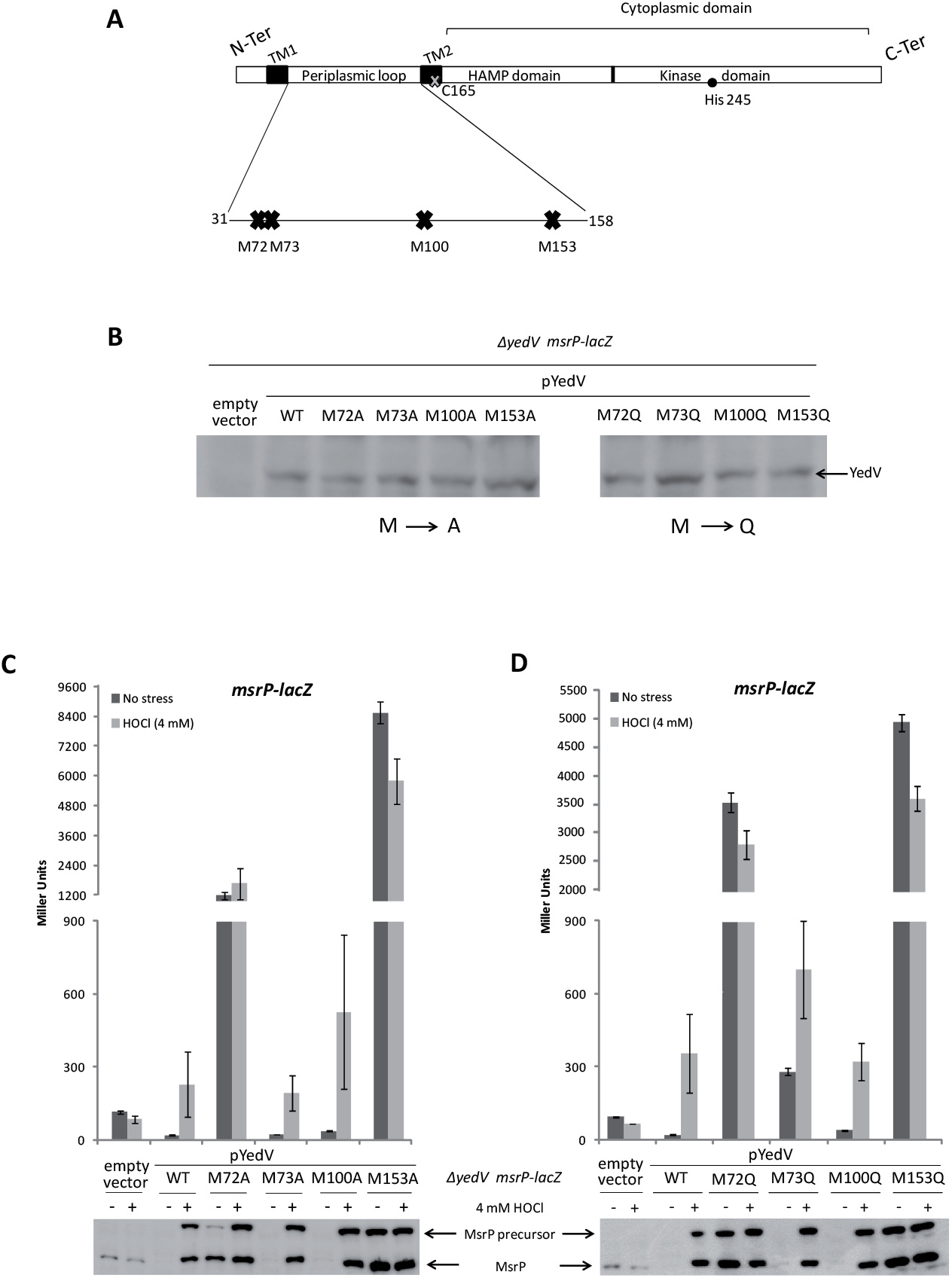
Role of each individual Met residues of the periplasmic loop of YedV. **A)** Schematic view of YedV sub-domains. YedV is a trans-membrane protein comprised of a small cytosolic N-Ter domain, two trans-membrane (TM) helices bordering a periplasmic domain. The periplasmic domain contains 4 Met residues and the second TM (TM2) contains the Cys165 pointed out by *Urano et al*.^16^as being important for YedV. The C-ter cytosolic domain is formed by the HAMP domain and by the catalytic domain harboring the kinase activity. **B)** Evaluation of the expression of different plasmidic alleles of *yedV* in strain CH186. CH186 cells carrying the empty plasmid pBBRMCS1 or expressing a variant of yedV (Wilde type or mutants) were cultured over-night at 37 °C in LB medium in presence of 1 mM IPTG. Immunoblot analysis were carried out using rabbit anti-YedV as a primary antibody. **C-D)** Effect of single Met mutations on *msrP* expression. The expression of *msrP* is used as a read out of YedVW activity. MsrP production is assessed by immunoblot analysis using anti-MsrP as a primary antibody and by measuring *msrP-lacZ* expression. Both analyses were carried out on CH186 cells carrying either the empty plasmid pBBRMCS1 (negative control), expressing wild-type *yedV* (positive control) or *yedV* mutants. *msrP* expression was analyzed in each strain after treatment with 4 mM HOCl for 1 hour in LB medium. Values of the ß-galactosidase assays are expressed as means ± SD (n=3). **C)** Effect of Met-to-Ala substitution on YedV activity: M72A, M73A, M100A, M153A **D)** Effect of Met-to-Gln substitution on YedV activity: M72Q, M73Q, M100Q, M153Q.

We reasoned that the variant containing Met-to-Ala would inform us on the physicochemical constraints prevailing at the position, and the variant containing Gln, a mimetic of Met-O^18^, would inform us specifically on the effect of oxidizing each Met on YedV activity. To test the effect of these substitutions, a *yedV*-less host carrying the *msrP-lacZ* fusion (strain *CH186*) was complemented with plasmids expressing mutated variants of *yedV*, as well as the plasmid carrying the WT gene. All the YedV variant proteins exhibit a similar level of expression as revealed by western-blot analysis (Fig. 4B). Next, the strains were grown and treated with HOCl (4 mM) for 1 hour. The *msrP-lacZ* activity and the production of MsrP were monitored. As already reported, a *yedV*-null mutant has a higher basal expression of *msrP-lacZ* compared to the strain expressing the WT gene, and it cannot respond to HOCl^13^ (Fig. 4C). These phenotypes can be complemented with a plasmidic *yedV*^*WT*^ allele: first, we found low protein levels of MsrP and a decrease in β-galactosidase activity of the *msrP-lacZ* fusion in the absence of HOCl. Then, we observed that the presence of HOCl led to a high steady-state level of MsrP and an increase of the β-galactosidase activity from 20 to 230 Miller Units (Fig. 4C). We next assayed *msrP* expression in the strain *CH186* complemented with different YedV mutants with or without HOCl. The mutation of residue Met 100 into either Ala or Gln had no drastic effect on YedV’s activity. These mutants show similar levels of β-galactosidase activity and MsrP production compared to YedV^WT^ in the absence and in the presence of HOCl (Fig. 4C, Fig. 4D). The mutation of residue Met73 into Ala had no effect on YedV activity either (Fig. 4C). However, when residue Met73 was mutated into a Met-O mimicking residue (Gln) we observed a higher basal expression of *msrP* compared to YedV^WT^ in the absence of HOCl (around 20 and 280 Miller Units with YedV^WT^ and YedV^M73Q^ respectively). When HOCl was added, we observed an increase in *msrP* expression and β-galactosidase activity to higher levels than those observed with the WT allele (around 350 and 700 Miller Units with YedV^WT^ and YedV^M73Q^ respectively) (Fig. 4D). Mutating residues Met72 or Met153 into either Ala or Gln resulted in an unexpected observation: these mutations shift the sensor to a “locked-on” state. Strains expressing these variants show a high level of MsrP and exacerbated expression of *msrP-lacZ* even in the absence of HOCl (around 1,100 and 8,500 Miller Units with YedV^M72A^ and YedV^M153A^ and 3500 and 4900 Miller Units with YedV^M72Q^ and YedV^M153Q^ respectively). The β-galactosidase activity is not affected by the addition of HOCl and is maintained at high levels (around 1,600 and 5,800 Miller Units with YedV^M72A^ and YedV^M153A^ and 2800 and 3600 with YedV^M72Q^ and YedV^M153Q^ respectively) (Fig. 4C, Fig. 4D). Taken together, these observations suggest that Met72 and Met153 are important for YedV activity.

To further investigate the roles of Met72 and Met153, we have constructed mutants of YedV, each carrying a combination of two, three or four Met-to-Ala substitutions. These variants proteins exhibit a similar level of expression as revealed by western-blot analysis (Fig. 5A-C). These mutants were introduced into strain *CH186*. The *msrP-lacZ* activity and the production of MsrP were monitored in the presence and absence of 4 mM HOCl. We observed that mutating simultaneously Met73 and Met100 into Ala had no effect on YedV activity. Similar levels of β-galactosidase activity and MsrP production are observed compared to YedV^WT^ in the absence and in the presence of HOCl, showing on more that they are not essential for YedV activity (Fig. 5B). When we combined mutation M73A to M72A, we observed a decrease in the basal expression of *msrP* compared to that observed with YedV^M72A^ (1175 and 459 Miller Unit with YedV^M72A^ and YedV^M72-73A^ respectively). The addition of HOCl increased the expression of *msrP* in the presence of YedV^M72-73A^ whereas no increase in *msrP* expression was observed with YedV^M72A^ after the addition of HOCl (Fig. 5B). YedV^M72-100A^ shows the same phenotype than the variant YedV^M72A^: same high basal level of *msrP* expression in the absence of HOCl (around 1180 and 1300 Miller Unit with YedV^M72A^ and YedV^M72-100A^ respectively). The addition of HOCl did not affect the expression of *msrP* and it was maintained approximately at the same level in both mutants (around 1600 and 1900 Miller Unit with YedV^M72A^ and YedV^M72-100A^ respectively). These results show that M72A is a dominant mutation compared to M100A (Fig. 5B). Mutations M72A or M73A seem to be recessive compared to mutation M153A. The variants YedV^M72-153A^ and YedV^M73-153A^ show very high levels of *msrP-lacZ* expression those are comparable to those observed with the YedV^M153A^ variant (around 8550, 5550 and 6450 Miller Units with YedV^M153A^, YedV^M72-153A^ and YedV^M73-153A^ respectively). The addition of HOCl does not affect the expression of *msrP* (around 5800, 5100 and 4760 Miller Units with YedV^M153A^, YedV^M72-153A^ and YedV^M73-153A^ respectively) (Fig. 5B). To assess the importance of residue Met153 in YedV activity, we have constructed a YedV mutant carrying the simultaneous substitution of Met72, Met73 and Met100 into Ala. This way, only Met153 is available to react with HOCl within the periplasmic loop. This mutant was not able to complement *yedV* deletion in strain *CH186*: the basal level of *msrP-lacZ* is around 608 Miller Unit in absence of HOCl. When HOCl is added, *msrP-lacZ* expression increases up to around 1000 Miller Units. Nevertheless, when we compared the fold-induction between YedV^M72-73-100A^ and YedV^WT^ in response to HOCl, we find a 30-fold increase of *msrP-lacZ* in a YedV^WT^ background and only a two-fold increase in a YedV^M72-73-100A^ background. This observation suggests that the presence of Met153 alone is not sufficient for a fully functional YedV (Fig. 5C). Based on these observations, we hypothesized that all Met residues might be important for YedV activity and that substituting all Met residue into Ala might render YedV blind to the signal. Nevertheless, the phenotype observed with YedV^M72-73-100-153A^ is comparable to that observed with YedV^M153A^: the variant has a “locked on state” and the addition of HOCl does not affect the expression of *msrP-lac*Z that is maintained at high levels. This phenotype might likely be due to the dominant mutation M153A (Fig. 5C).

**Figure 5-.**
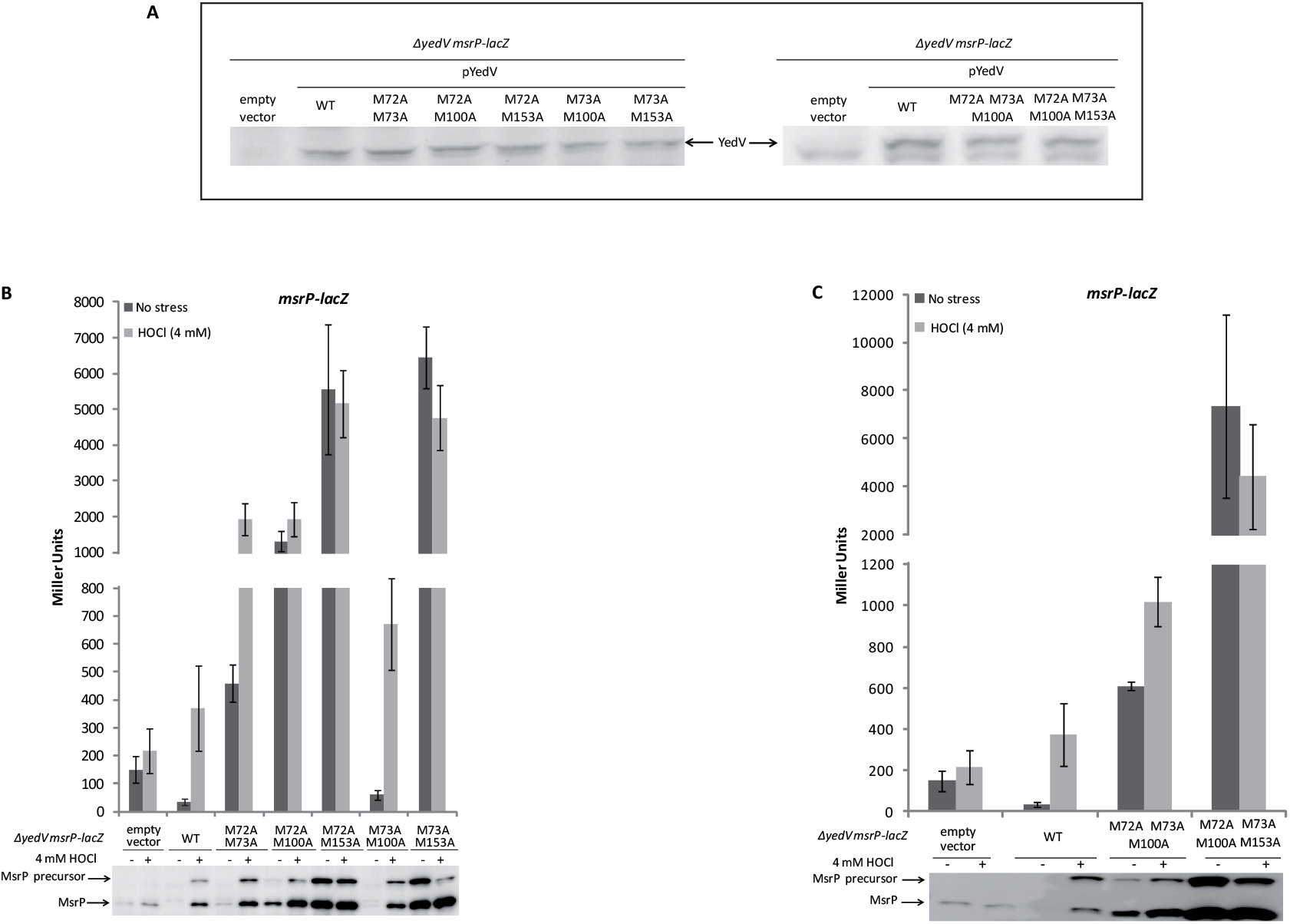
Effect of combining two, three or four Met-to-Ala mutations on YedV activity. **A)** Immunoblot analysis showing the expression of different plasmidic *yedV* alleles (WT and double Ala-to Met mutations) in CH186 cells. CH186 cells carrying the empty plasmid pBBRMCS1 or expressing a variant of yedV (WT or mutants) were cultured over-night at 37 °C in LB medium in presence of 1 mM IPTG. Immunoblot analysis were carried out using rabbit anti-YedV as a primary antibody. **B)** Effect of combining two Met-to-Ala mutations on YedV activity. The expression of *msrP* is used as a read out of YedVW activity. MsrP production is assessed by immunoblot analysis using anti-MsrP antibody and by measuring *msrP-lacZ* expression. Both analysis were carried out on CH186 cells carrying either the empty plasmid pBBRMCS1 (negative control), expressing wild-type *yedV* (positive control) or *yedV* mutants: M72-73A, M72-100A, M72-153A, M73-100A, M73-153A. *msrP* expression was analyzed in each strain after treatment with HOCl. Values of the ß-galactosidase assays are expressed as mean ± SD (n =3). **C)** Effect of combining three or four Met-to-Ala mutations on YedV activity. The expression of *msrP* is used as a read out of YedVW activity. MsrP production is assessed by immunoblot analysis using anti-MsrP antibody and by measuring *msrP-lacZ* expression. Both analysis were carried out on CH186 cells carrying either the empty plasmid pBBRMCS1 (negative control), expressing wild-type *yedV* (positive control) or the *yedV* mutants: M72-73-100A or M72-73-100-153A. *msrP* expression was analyzed in each strain after treatment with 4 mM HOCl. Values of the ß-galactosidase assays are expressed as mean ± SD (n =3).

### Negative feedback of MsrP on the expression of the hiuH-msrP-msrQ operon

The Met redox switch YedV-activation hypothesis implies that deletion of the *msrP* gene should modify the pattern of HOCl-YedVW dependent induction of *hiuH-lacZ* expression as MsrP would be able to reduce Met-O containing YedV to turn off the activation once the HOCl signal is gone. To test this hypothesis, we have constructed the strains SH100 (MsrP^+^) and SH101 (MsrP^-^) in which the *yedWV* operon was under the control of the tetracycline resistance (TetR) promoter, in order to abolished HOCl-dependent *de novo* synthesis of the sensor. Both strains were grown in minimum medium. HOCl was added at OD = 0.2 and the β-galactosidase activity (*hiuH-lacZ)* was monitored over time. Our results show that HOCl treatment leads to a higher level of β-galactosidase activity in the SH101 MsrP^-^) compared to the SH100 (MsrP^+^). No difference was observed in absence of HOCl (Fig. 6A). Moreover, the over-production of MsrP using a plasmid led to a significant decrease of the induction of *hiuH-lacZ* during HOCl stress (Fig. 6B). Together, these results show a negative feedback of MsrP on YedW-regulated genes (i.e. *hiuH*) during HOCl stress, most likely involving a methionine redox control process.

**Figure 6-.**
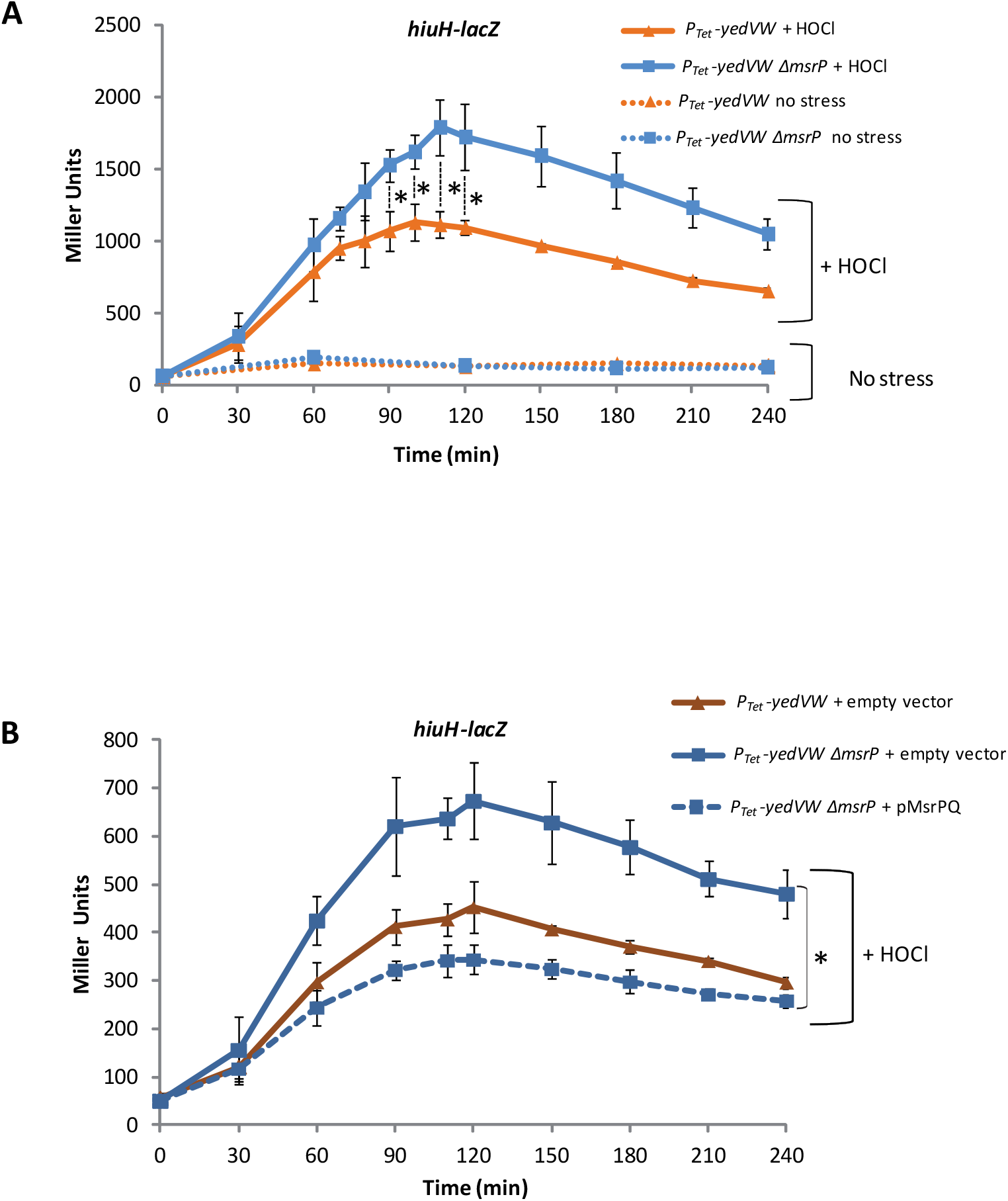
Effect of MsrP on *hiuH-lacZ* expression. **A)** Effect of *msrP* deletion on *hiuH-lacZ* expression. Strains SH100 (*P*_*TetR*_*-YedVW cat-hiuH-lacZ*) and SH101 (*P*_*TetR*_*-YedVW ΔmsrP cat-hiuH-lacZ*) were cultured in minimal medium M9. 135 µM of HOCl was added to each strain at OD_600_=0.2, and *hiuH-lacZ* expression was followed over 4 hours. Values of the ß-galactosidase assays are expressed as means ± SD (n=4). An asterisk indicates statistically significant difference between ß-galactosidase activity of different strains. **B)** Effect of complementing *msrP* deletion on *hiuH-lacZ* expression. SH100 carrying an empty plasmid and SH101 carrying either an empty plasmid or pMsrPQ were cultured in minimal medium M9. 135 µM HOCl were added to each strain at OD_600_=0.2 and *hiuH-lacZ* expression was followed over 4 hours. Values of the ß-galactosidase assays are expressed as means ± SD (n=3). An asterisk indicates statistically significant difference between ß-galactosidase activity of different strains. *P ≤ 0.05 (Mann-Whitney U test).

### Residues Met72 and Met153 are highly conserved in YedV homologues

In order to support a functional role for the periplasmic Met residues in YedV, we assessed their conservation in YedV sequence. To do so, we first established the list of YedV homologues from the STRING database. We defined as homologue, the two-component systems that have the same genetic environment as YedV of *E. coli*, e.g. next to an *hiuH* or *msrP*-*msrQ* genes. Based on this criterion, we selected 12 TCS including *E. coli* YedV (Fig. 7A). 10 of the selected homologues were found in Gammaproteobacteria and only 2 were found in Betaproteobacteria (*Herbaspirillum frisingense* and *Herminiimonas arsenicoxydans*). This indicates that *yedWV* might not be widely distributed among prokaryotes. We next aligned the protein sequence of the different homologues of YedV. We found that Met72 and Met153 are highly conserved: Met72 is found in 92 % of the homologues (11 out of the 12 analyzed sequence) and Met153 is conserved in 75% of the homologues (9 out of the 12 homologues). Conversely, Met73 is less conserved, being only found in 58% of the homologues (7 out of 12). Met100 is poorly conserved and occurs only in *Shigella flexneri* and *E. coli* YedV (Fig. 6B). Taken together, this analysis suggest that Met72 and Met153 might be important residues for the sensor YedV.

**Figure 7-.**
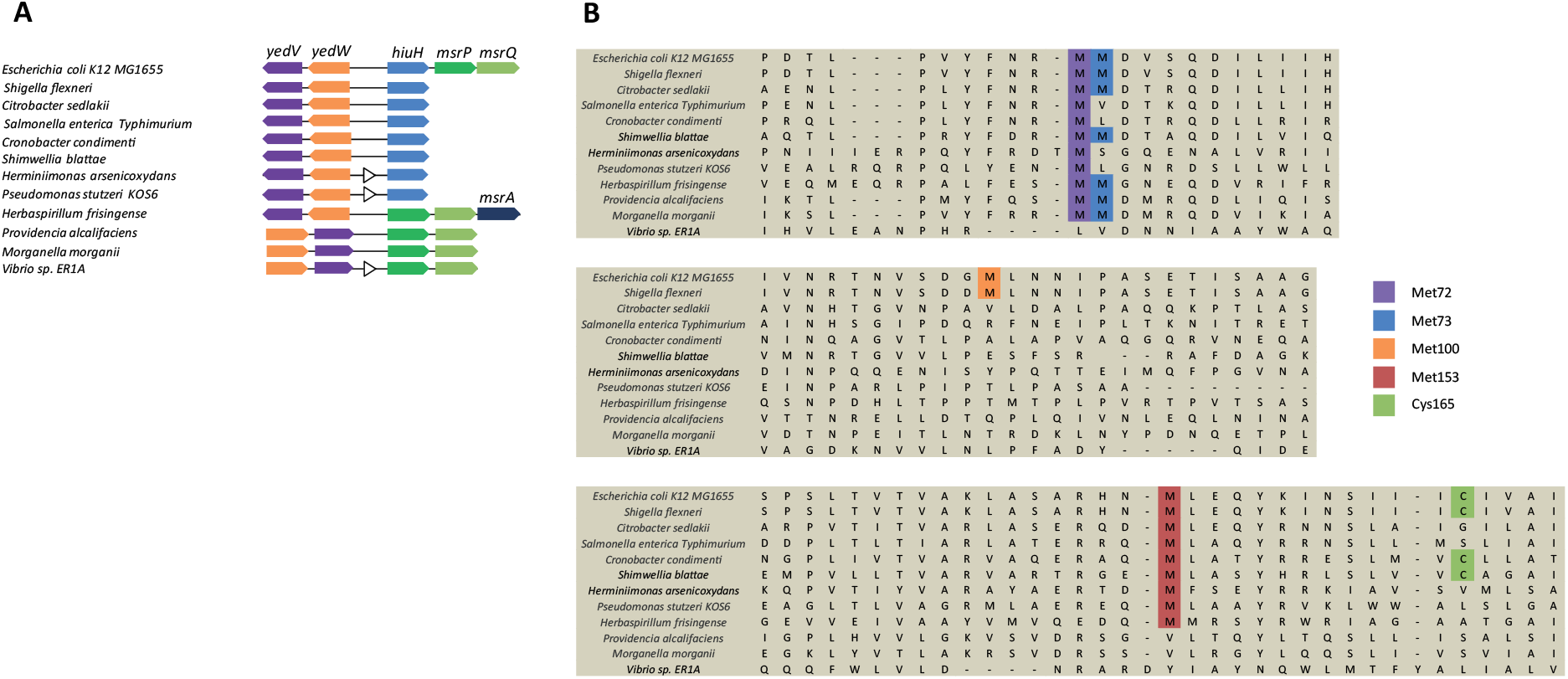
Protein sequence comparison between YedV homologues. YedV Homologues were selected based on their genetic environment (close to an *msrP-msrQ gene* or an *hiuH* gene) and on their orthology relation with *E. coli yedV*. **A)** Scheme representing the genetic environment of each selected YedV homologue. **B)** Protein sequence comparison between different YedV homologues. The alignment shown in this figure represents three different regions of YedV each containing one or two residue(s) of interest: Met72, Met73, Met100, Met153 and Cys165. Sequence comparison was carried-out using Clustal Omega (Whitworth and Cock 2009). The code attributed to each homologue in the STRING database is the following: *b1968* (*Escherichia coli K12 MG1655*), *SF2015* (*Shigella flexneri*), *BBNB01000011_gene586* (*Citrobacter sedlakii*), *CY43_05595* (*Salmonella enterica Typhimurium*), *BN137_3921* (*Cronobacter condimenti*), *EBL_c13160* (*Shimwellia blattae*), *HEAR0605* (*Herminiimonas arsenicoxydans*), *B597_003020* (*Pseudomonas stutzeri KOS6*), *HFRIS_015940* (*Herbaspirillum frisingense*), *PROVALCAL_02806* (*Providencia alcalifaciens*), *MU9_2071* (*Morganella morganii*), *HW45_14750* (*Vibrio sp. ER1A*).

### The trans-membrane residue Cys165 is important for signal transduction

Previous mutagenesis approach, using a Cys-to-Ala substitution, highlighted the residue Cys165, located in the second trans-membrane domain (TM2), as important for YedV activation by H_2_O_2_^16^. We therefore decided to investigate the role of Cys165 in YedV activation during HOCl stress. First, the C165A mutant was constructed and we observed a similar abundance of YedV production compared to the wild type strain (Fig. 8A). Next, we showed that in presence of HOCl, the expression of the *msrP-lacZ* fusion and the MsrP protein levels were lower in strains expressing YedV^C165A^ compared to the wild type (Fig. 8B). However, this residue was shown to be only conserved in 4 YedV homologues, challenging its functional role in YedV activity (Fig. 7B). We next considered the possibility that Cys165 might contribute to the transmission and not to the sensing of the signal. To investigate this hypothesis, we took advantage of the M72Q variant, which caused constitutive activation of YedV, to assess this question. We built the M72Q-C165A mutant, which exhibits a level of YedV production similar to the WT (Fig. 8A). By measuring the expression of the *msrP-lacZ* fusion and the MsrP protein levels, we showed that the strain expressing YedV^M72Q-C165A^ has a lower MsrP level than the strain expressing YedV^M72Q^ (Fig. 8B). Taken together, these results show that C165A mutation was dominant on M72Q and suggest that Cys165 is an important residue for signal transduction from the periplasmic sensing domain to the cytosolic kinase domain.

**Figure 8-.**
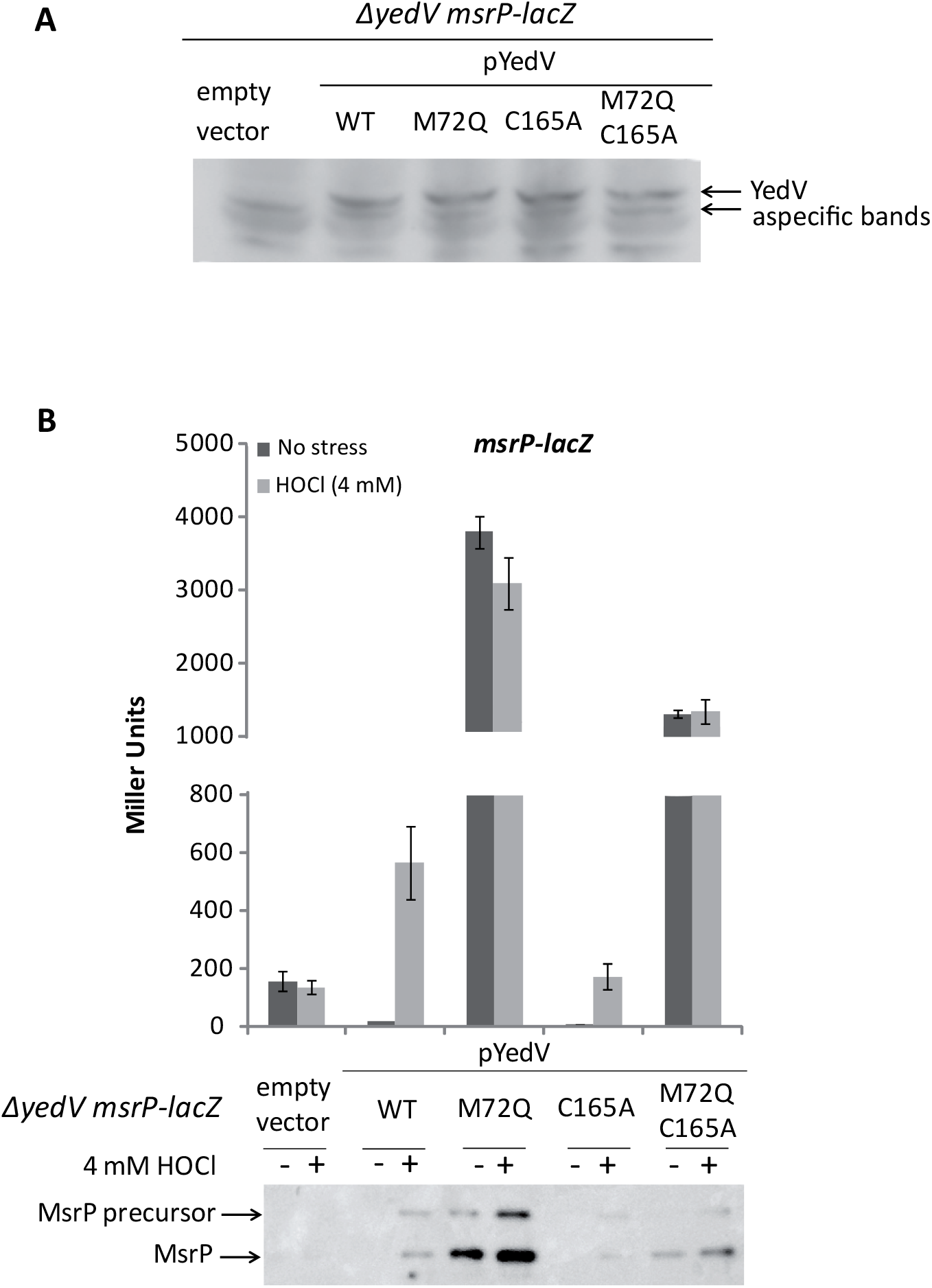
Evaluation of the role of Cys165 in YedV activity. **A)** Immunoblot analysis showing the expression of different plasmidic *yedV* alleles: WT, M72Q, C165A and M72Q-C165A in CH186 cells. CH186 cells carrying the empty plasmid pBBRMCS1 or expressing a variant of *yedV* (WT or mutants) were cultured over-night at 37 °C in LB medium in presence of 1 mM IPTG. Immunoblot analysis were carried out using rabbit anti-YedV as a primary antibody. **B)** Effect of Cys165A mutation on YedV activity. The expression of *msrP* is used as a read out of YedVW activity. MsrP production is assessed by immunoblot analysis using anti-MsrP antibody and by measuring *msrP-lacZ* expression. Both analysis were carried out on CH186 cells carrying either the empty plasmid pBBRMCS1 (negative control), expressing wild-type *yedV* (positive control) or *yedV* mutants: M72Q, C165A or M72Q-C165A. *msrP* expression was analyzed in each strain after treatment with 4 mM HOCl for 1 hour in LB medium. Values of the ß-galactosidase assays are expressed as mean ± SD (n =3).

## DISCUSSION

In bacteria, the two component systems (TCS) are widespread signaling pathways allowing cells to sense, respond and adapt to fluctuating environments and challenging conditions. Based on genome sequencing, *E. coli* MG1655 has been predicted to carry a total of 24 “classical” TCS composed of a histidine kinase and a response regulator ^19^. Many of them have yet to be characterized. Other well-studied TCS were shown to regulate bacterial metabolism, nutrient acquisition and stress response to numerous stimuli such as pH, metals and envelope integrity ^20,21^. Therefore, the characterization of *E. coli* TCS appears to be important for developing a complete knowledge of bacterial regulatory networks. The aim of this study was to further characterize the TCS YedVW in *E. coli* and we have found that oxidative stress mediated specifically by RCS (HOCl and *N*-ChT) can activate this system. We have also brought evidences concerning the molecular mechanism of signal sensing by YedV.

Until now, YedVW was poorly characterized and the first evidence for the YedVW-dependent regulon was brought by using SELEX methodology^15^. Four YedW binding sites were identified in the whole genome: one between the two operons *yedWV* and *hiuH-msrPQ*, one within the *cyoABCDE* operon encoding the cytochrome oxidase, one near the *qorA* gene encoding a quinone oxido-reductase and one between the *cusRS* and the *cusCFBA* operons, encoding the copper responding TCS and the copper efflux pump, respectively ^15^. However, this approach cannot exclude the possibility that some targeted genes could be regulated only by YedW and not by YedV via a cross-talk between TCS. Indeed, such a cross-talk have to be taken into consideration as YedW was shown to be phosphorylated by CusS, the copper-sensing HK ^20^. Thereby, previous results and those presented in this study, show that *hiuH-msrPQ* operon is regulated by the YedVW TCS ^13,15^.

As a first step in characterizing YedVW, we tried to identify the signals sensed by the HK YedV. Previous studies reported contradictory results. It has been shown that YedVW, in response to H_2_O_2_ stress, increases the mRNA level of its target gene *hiuH*, suggesting that YedV senses H_2_O_2_ ^15^. Another study conducted in our lab showed that YedVW induces the expression of the gene encoding the periplasmic methionine sulfoxide reductase system (MsrPQ) in response to HOCl and not to H_2_O_2_ ^13^. This conclusion was obtained at both transcriptional and translational levels, using an *msrP* reporter fusion and by assessing the MsrP protein levels, respectively ^13^. These contradictory results prompted us to re-investigate the signals sensed by YedV. In this study, we assessed both mRNA and protein levels of YedVW target genes. The strength of this study is the simultaneous comparison between the effect of HOCl and H_2_O_2_ on YedVW and the evaluation of the expression of target genes both at the transcriptional and translational level. Our results show that HOCl is a more physiological signal for YedVW compared to H_2_O_2_ and that the previously described activation of YedVW by H_2_O_2_ might be irrelevant and non-physiological as only a slight increase in mRNA levels is detected. We also tested other ROS/RCS that might be sensed by YedV. To do that, we treated cells with different compounds known to generate oxidative stress. Among all the compounds tested, we have only detected an activation of YedVW in response to HOCl and *N*-ChT. Our results suggest that YedV is a specific RCS sensor in *E. coli*. Based on these findings, we proposed to rename this system, HypVW in reference to hypochlorous acid. Until now, two different names were attributed to YedVW: HprSR for hydrogen peroxide sensor ^16^ and regulator and MsrVW ^22^ for its association with MsrPQ. We think that both names are not suitable for this TCS with light of the present study: first, we have shown, throughout this study, that H_2_O_2_ is not an adequate signal for YedVW. Second, the Msr denomination is misleading and should be exclusive for proteins that carry a methionine sulfoxide reductase enzymatic activity or that are essential for an Msr activity (such as MsrQ). Moreover and unfortunately, HypS and HypR are already attributed to other orthologs transcriptional regulators that are specific to HOCl in *Mycobacterium smegmatis* and in *Staphylococcus aureus* respectively ^23,24^. Therefore, HypVW appears to be a good compromise for the nomenclature.

As a second step, we investigated the molecular mechanism underlying HOCl detection by HypV. It has been shown that HOCl activates RCS-specific transcription factors through post-translational modification of Cys or Met residues. Indeed, HOCl activates RclR and NemR by a thiol-disulfide switch mechanism or by the formation of a Cys-Lys sulfenamide bonds, respectively, while HOCl oxidizes three Met residues in HypT, leading to its activation^9,8,10^. Interestingly, HypV contains four Cys and six Met residues. Two Cys residues (Cys165 and Cys172) are located in the second trans-membrane domain and the other two (Cys333 and Cys338) are in the cytoplasmic domain. In the case of Met residues, four of them are located in the periplasmic loop of HypV (Met72, Met73, Met100 and Met153), and the other two are in the cytoplasmic domain (Met225 and Met352). The involvement of Cys residues in HypV sensing mechanism has been already investigated and pointed-out to the trans-membrane Cys165 residue as being important for H_2_O_2_ detection ^16^. Met residues are also highly sensitive to HOCl and because usually HK uses their periplasmic loop to sense extracellular signals ^25^, we therefore focused our study on the 4 Met residues located in the periplasmic domain. We conducted a systematic Met-to-Ala and Met-to-Gln substitution of each Met residue. We also generated HypV mutants containing different combination of two, three or four Met-to-Ala substitutions. Replacing Met with Ala will prevent the sulfur atom to react with HOCl and substituting Met with Gln mimics Met-O residues ^26^. We observed that substituting Met72 or Met153 with Ala or Gln had drastic consequences on HypV, leading to a constitutive expression of *msrP*. Substituting Met73 with Gln residue led to a higher basal activity compared to HypV^WT^ but this variant was not blind to the signal. Mutations of residue Met100 (M100A and M100Q) were neutral and had no effect on HypV. The analysis of HypV double mutants show that M72A and M153A are dominant compared to M73A and M100A. We also showed that residues M100 and M73 are dispensable for HypV activation by HOCl. When analyzing mutant HypV^M72-73-100A^, it was clear that residue M153A alone cannot activate HypV to its full potential in response to HOCl, showing that other residues are implicated. We were not able to conclude on the implication of Met residues in HypV activity since the mutant HypV^M72-73-100-153A^ was constitutively active, likely due to M153A mutation. Altogether, these results show that Met72, 73 and 153 are important for HypV activity with a functional hierarchy based on the residues. Met72 and Met153 appear to be crucial for HypV whereas Met73 seem to have a secondary role. Strikingly, these conclusions somehow correlate with the fact that M72 and M153 are conserved throughout the HypV homologues whereas M73 is less conserved.

In addition to the mutational approach discussed earlier, we assayed the effect of MsrP on HypV activation. Reasoning that if the activation of HypV is linked to the oxidation of Met residues, then MsrP could participate in switching off HypV by reducing back the Met-O residues. Our results show that the level of *msrP-lacZ* expression is dependent on the level of MsrP production. Because the HOCl-dependent induction of *msrP-lacZ* is regulated by HypV, this observation strengthens a central role of the Met redox state in HypV activity.

Finally, we reinvestigated the role of the trans-membrane Cys165 residue using the HypV^M72Q^ constitutive variant. First, we confirm that an HypV^C165A^ variant is affected in its activity in agreement with the previous study ^16^. However, the localization and the lack of conservation of Cys165 lead us to postulate that this position (165) rather than the Cys residue itself (not conserved) could be important for signal transduction to the catalytic domain and not for signal sensing. The fact that C165A has been found to be dominant on the M72Q mutation, even in the absence of the signal, comfort this hypothesis.

Based on all the findings cited above, it appears that a Met-redox switch might play a role in HypV sensing mechanism, in sharp homology with the HOCl-specific transcription factor HypT. This means that the oxido-reduction state of particular Met residues (we speculate Met72 and Met153) might participate in the activation of HypV by HOCl. Also, MsrP might play a role in shutting down the system by reducing the oxidized Met residues, forming a negative feedback loop.

If Met residues are implicated in HypV activation, this might explain why H_2_O_2_ cannot activate properly HypV: in terms of kinetics, the oxidizing reactivity of Met residues with H_2_O_2_ is much lower than with HOCl (2 × 10^2^ M^−1^·s ^−1^ for H_2_O_2_ versus 3.4 × 10^7^ M^−1^·s ^−1^ for HOCl) ^17^ making the reaction much slower and requiring higher concentration of H_2_O_2_ that might not be physiological. One should note that we didn’t bring direct proof of *in vivo* Met oxidation in HypV. Therefore additional studies will be required to consolidate our proposed model. Also, our data do not exclude the possibility that HypV might be activated indirectly by RCS through a Met-oxidation process of a third periplasmic component which might interact with HypV. Our model needs to be consolidated in the future by a biochemical approach to assess the different post-translational modification undergone in HypV during RCS stress in *vivo*.

Taken together, we provide strong evidence that *E. coli* acquired a two-component system that detects and respond to RCS molecules and to our knowledge, HypVW is the only RCS related TCS in *E. coli*. Unlike ROS, information about RCS response in bacteria is still poorly understood. The further characterization of HypVW is an important issue for understanding RCS killing and resistance in prokaryotes.

## MATERIAL AND METHODS

### Strains and microbial techniques

The strains used in this study are listed in table 1. To construct strain SH198 (*yedW* knock-out strain), the *yedW* mutant allele from the Keio collection ^27^ was transferred to *E. coli* MG1655 strain (wild-type) using P1 transduction standard procedure ^28^. Kanamycin resistant strains were selected and verified by PCR. To excise the kanamycin cassette, we used pCP20 plasmid ^27,29^. The *hiuH-lacZ* fusion in strain CH184 was constructed using the protocol described by Mandin and Gottesman ^30^. First, the promoter region of *hiuH* located between nucleotide -274 and the ATG initiation triplet was amplified using the primers lacI-hiuH-For and lacZ-hiuH-Rev. Using mini lambda recombining, the PCR product was recombined in the chromosome of an *E. coli* strain carrying a P_BAD_-cat-SacB cassette inserted in front of the gene *lacZ* (strain PM1205). Recombinants who lost the cat-sacB cassette were selected and verified by PCR. Those clones carry the translational fusion *hiuH-lacZ*. Strain LL1700 containing a CmR cassette was constructed in order to transfer the *hiuH-lacZ* fusion into an *E. coli* MG1655 (Wilde-type) background. To do so, we inserted a resistance cassette close to the fusion enabling its transduction: a Cat cassette was amplified using primers 5deI and 3deI. The resulting product carry on its 5’ end, a 40 bp homologous sequence to *lacI* and on its 3’ end, a 41 bp homologous sequence to *hiuH* promoter (located between the nucleotide -133 and -92, before the YedW boxes). The amplicon was then electroporated into the strain CH184 carrying the pKD46 plasmid. Cat resistant clones were selected and tested by PCR. The *lacI-cat-hiuH-lacZ* construct was then transferred to MG1655 cells using P1 transduction standard procedure. Chloramphenicol resistant clones were selected and verifies by PCR. Strain SH100 was constructed as follows. First a TetR-Kan sequence composed of the promoter TetR and the kanamycine cassette was amplified using the plasmid pBbS2K as template and the primers YedW_TetR_F and Kan_Boxyedx_R. The resulting PCR product share a 48 bp homologous sequence with *yedW* and a 41 bp homologous sequence with the intergenic region between *yedW* and *hiuH* located before the YedW boxes. After purification, the PCR product was transformed by electroporation into *E. coli* BW25113 cells (WT) carrying the plasmid pKD46 ^29^. Kan resistant clones were selected and verified by PCR. This construction was then transferred into LL1700 cells expressing *hiuH-lacZ* fusion using P1 transduction standard procedures. Selected clones were verified by PCR. The same procedure was followed to construct strain SH101, only the TetR-Kan amplicon was electroporated into an *E. coli ΔmsrP* strain.

**Table 1-.**
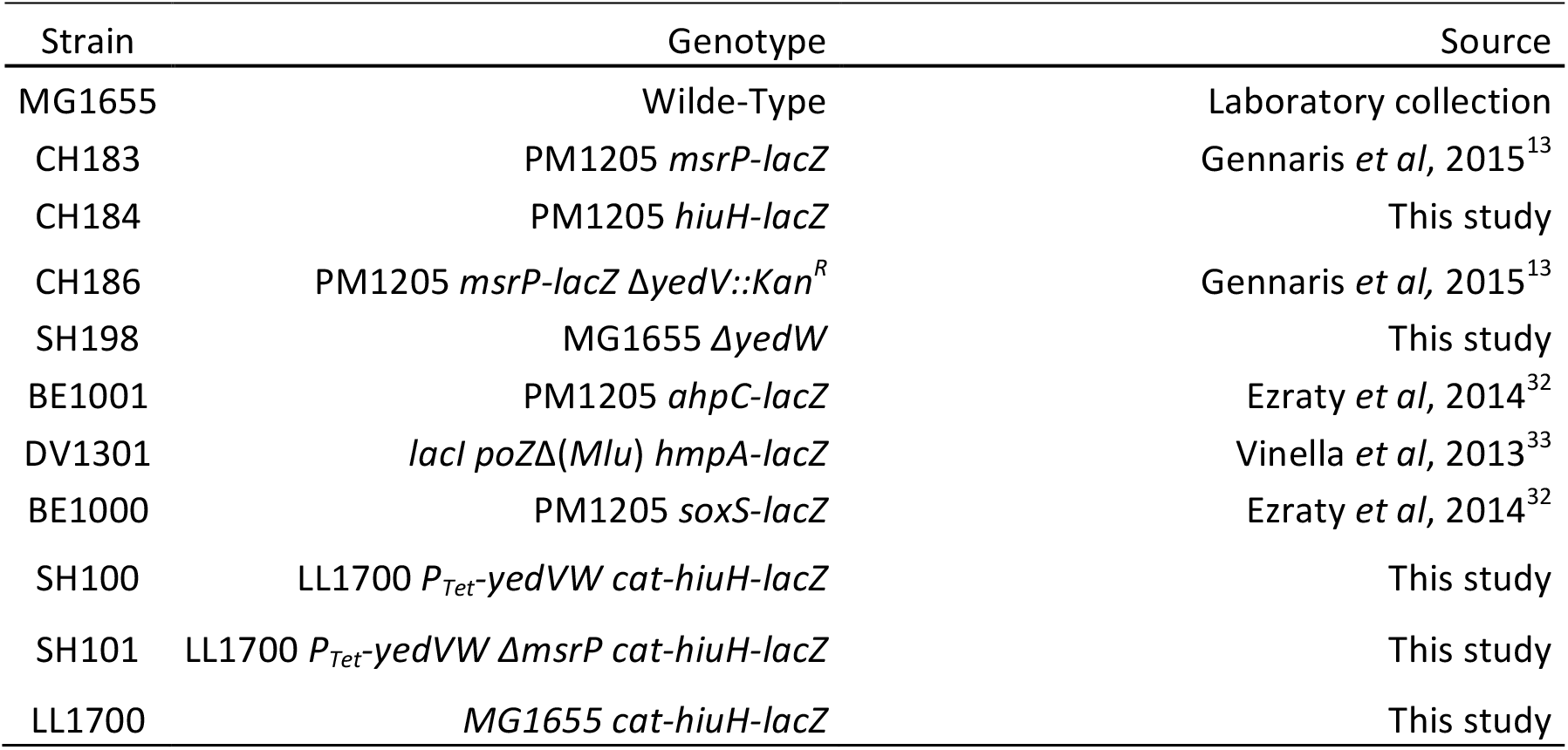
Strains used in this study. This table contains information regarding the strains used in this study, including their name, their genotype and their source.

| Strain | Genotype | Source |
| --- | --- | --- |
| MG1655 | Wilde-Type | Laboratory collection |
| CH183 | PM1205 *msrP-lacZ* | Gennaris *et al*, 2015<sup>13</sup> |
| CH184 | PM1205 *hiuH-lacZ* | This study |
| CH186 | PM1205 *msrP-lacZ ΔyedV::Kan<sup>R</sup>* | Gennaris *et al*, 2015<sup>13</sup> |
| SH198 | MG1655 *ΔyedW* | This study |
| BE1001 | PM1205 *ahpC-lacZ* | Ezraty *et al*, 2014<sup>32</sup> |
| DV1301 | *lacI poZΔ(Mlu) hmpA-lacZ* | Vinella *et al*, 2013<sup>33</sup> |
| BE1000 | PM1205 *soxS-lacZ* | Ezraty *et al*, 2014<sup>32</sup> |
| SH100 | LL1700 *P<sub>Tet</sub>-yedVW cat-hiuH-lacZ* | This study |
| SH101 | LL1700 *P<sub>Tet</sub>-yedVW ΔmsrP cat-hiuH-lacZ* | This study |
| LL1700 | MG1655 *cat-hiuH-lacZ* | This study |

### Plasmid constructions

The plasmids and primers used in this study are listed in table 2 and 3 respectively. The pYedV plasmid was constructed as follows. *yedV* was amplified from the chromosome of *E. coli* (MG1655) using primers XhoI_YedV_atg_F and HindIII_YedV_R. The PCR product was cloned into pBBRMCS1 plasmid using XhoI and HindIII restriction sites, generating plasmid pYedV. In this plasmid, the expression of YedV is under the control of the P_lac_ promoter inducible by IPTG. Plasmids pYedVM72A, pYedVM73A, pYedVM100A, pYedVM153A, pYedVM72Q, pYedVM73Q, pYedVM100Q, pYedVM153Q and pYedVC165A are derived from pYedV and carry single mutations of the *yedV* gene. Each mutant was constructed by site directed mutagenesis using two step PCR method: two overlapped PCR products were generated from pYedV using one external primer (either XhoI_YedV_atg_F or HindIII_YedV_R) and an internal primer carrying the desired mutation. After purification, the PCR products were used as templates for the second PCR carried out with the external primers only XhoI_YedV_atg_F and HindIII_YedV_R. The resultant PCR fragment was digested with XhoI and HindIII restriction enzymes (Biolabs) then cloned into pBBRMCS1. The same procedure was used to construct pYedVM72Q/C165 using pYedVM72Q as template and YedV_C165A_2_F and YedV_C165A_2_R as internal primers. *yedV* mutants carrying different combination of double Met-to-Ala substitutions were constructed using the two step PCR method described earlier: pYedV^M72-73A^, pYedV^M72-100A^, pYedV^M72-153A^, pYedV^M73-100A^, pYedV^M73-153A^. For that, pYedV plasmids carrying one of the two mutations was used as a template and the internal primers used for the PCR reaction carry each the second mutation to be introduced. All double mutants are cloned in pBBRMCS1. Mutant *yedV*^*M72-73-100A*^ was constructed using pYedV^M72-73A^ as template and YedV_M100A_F and YedV_M100A_R as internal primers. Mutant *yedV*^*M72-73-100-153A*^ was constructed using pYedV^M72-73-100A^ as template and YedV_M153A_F and YedV_M153A_R as internal primers. Both mutants *yedV*^*M72-73-100A*^ and *yedV*^*M72-73-100-153A*^ were cloned in pBBRMCS1. All PCR reactions were carried out using Phusion high fidelity Taq polymerase (Thermo Fisher).

**Table 2-.**
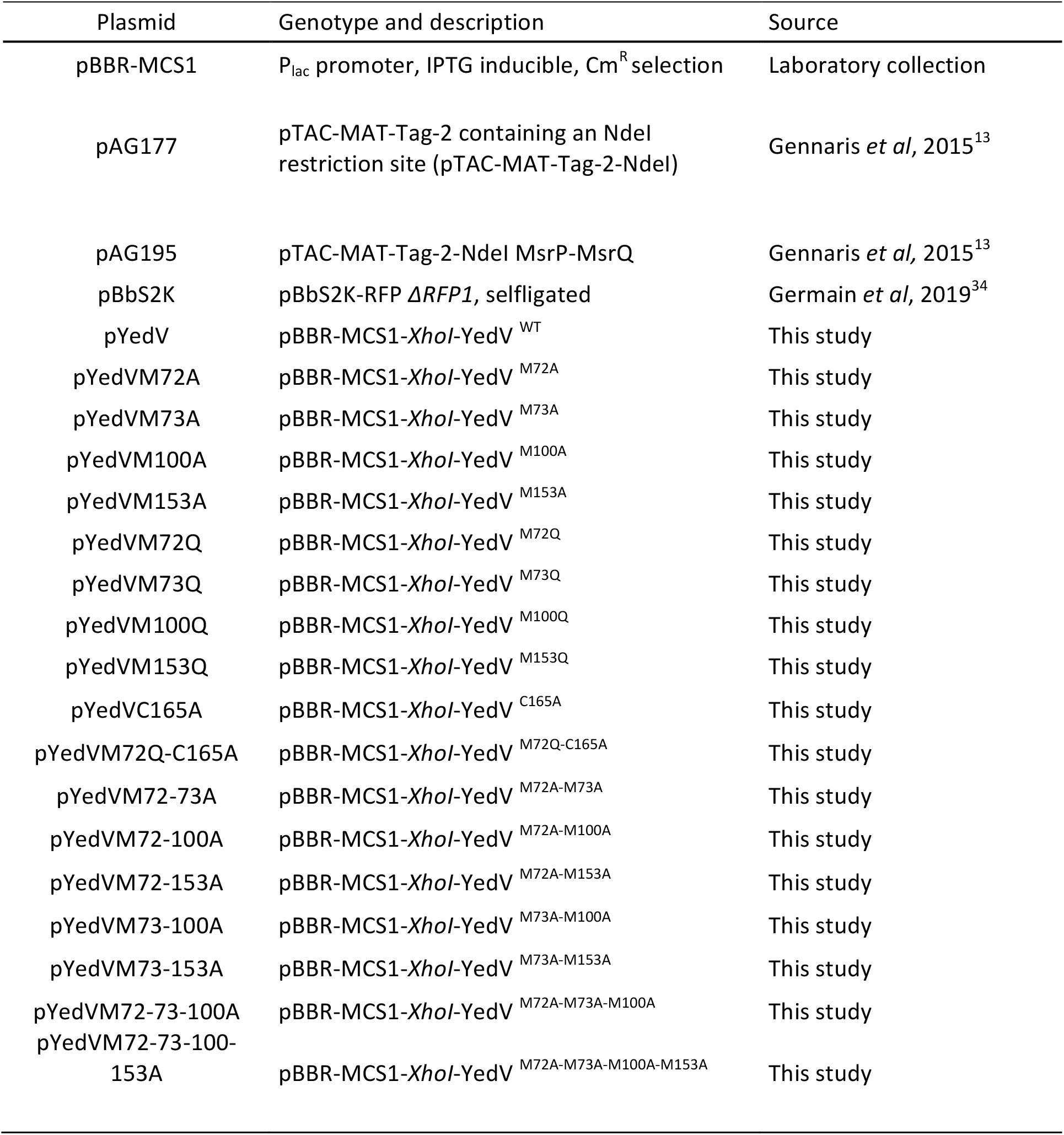
Plasmids used in this study. This table contains information regarding the plasmids used in this study, including their name, their genotype and their source.

| Plasmid | Genotype and description | Source |
| --- | --- | --- |
| pBBR-MCS1 | P<sub>lac</sub> promoter, IPTG inducible, Cm<sup>R</sup> selection | Laboratory collection |
| pAG177 | pTAC-MAT-Tag-2 containing an NdeI restriction site (pTAC-MAT-Tag-2-NdeI) | Gennaris *et al*, 2015<sup>13</sup> |
| pAG195 | pTAC-MAT-Tag-2-NdeI MsrP-MsrQ | Gennaris *et al*, 2015<sup>13</sup> |
| pBbS2K | pBbS2K-RFP $\Delta RFP1$, selfligated | Germain *et al*, 2019<sup>34</sup> |
| pYedV | pBBR-MCS1-*XhoI*-YedV<sup>WT</sup> | This study |
| pYedVM72A | pBBR-MCS1-*XhoI*-YedV<sup>M72A</sup> | This study |
| pYedVM73A | pBBR-MCS1-*XhoI*-YedV<sup>M73A</sup> | This study |
| pYedVM100A | pBBR-MCS1-*XhoI*-YedV<sup>M100A</sup> | This study |
| pYedVM153A | pBBR-MCS1-*XhoI*-YedV<sup>M153A</sup> | This study |
| pYedVM72Q | pBBR-MCS1-*XhoI*-YedV<sup>M72Q</sup> | This study |
| pYedVM73Q | pBBR-MCS1-*XhoI*-YedV<sup>M73Q</sup> | This study |
| pYedVM100Q | pBBR-MCS1-*XhoI*-YedV<sup>M100Q</sup> | This study |
| pYedVM153Q | pBBR-MCS1-*XhoI*-YedV<sup>M153Q</sup> | This study |
| pYedVC165A | pBBR-MCS1-*XhoI*-YedV<sup>C165A</sup> | This study |
| pYedVM72Q-C165A | pBBR-MCS1-*XhoI*-YedV<sup>M72Q-C165A</sup> | This study |
| pYedVM72-73A | pBBR-MCS1-*XhoI*-YedV<sup>M72A-M73A</sup> | This study |
| pYedVM72-100A | pBBR-MCS1-*XhoI*-YedV<sup>M72A-M100A</sup> | This study |
| pYedVM72-153A | pBBR-MCS1-*XhoI*-YedV<sup>M72A-M153A</sup> | This study |
| pYedVM73-100A | pBBR-MCS1-*XhoI*-YedV<sup>M73A-M100A</sup> | This study |
| pYedVM73-153A | pBBR-MCS1-*XhoI*-YedV<sup>M73A-M153A</sup> | This study |
| pYedVM72-73-100A | pBBR-MCS1-*XhoI*-YedV<sup>M72A-M73A-M100A</sup> | This study |
| pYedVM72-73-100-153A | pBBR-MCS1-*XhoI*-YedV<sup>M72A-M73A-M100A-M153A</sup> | This study |

**Table 3-.**
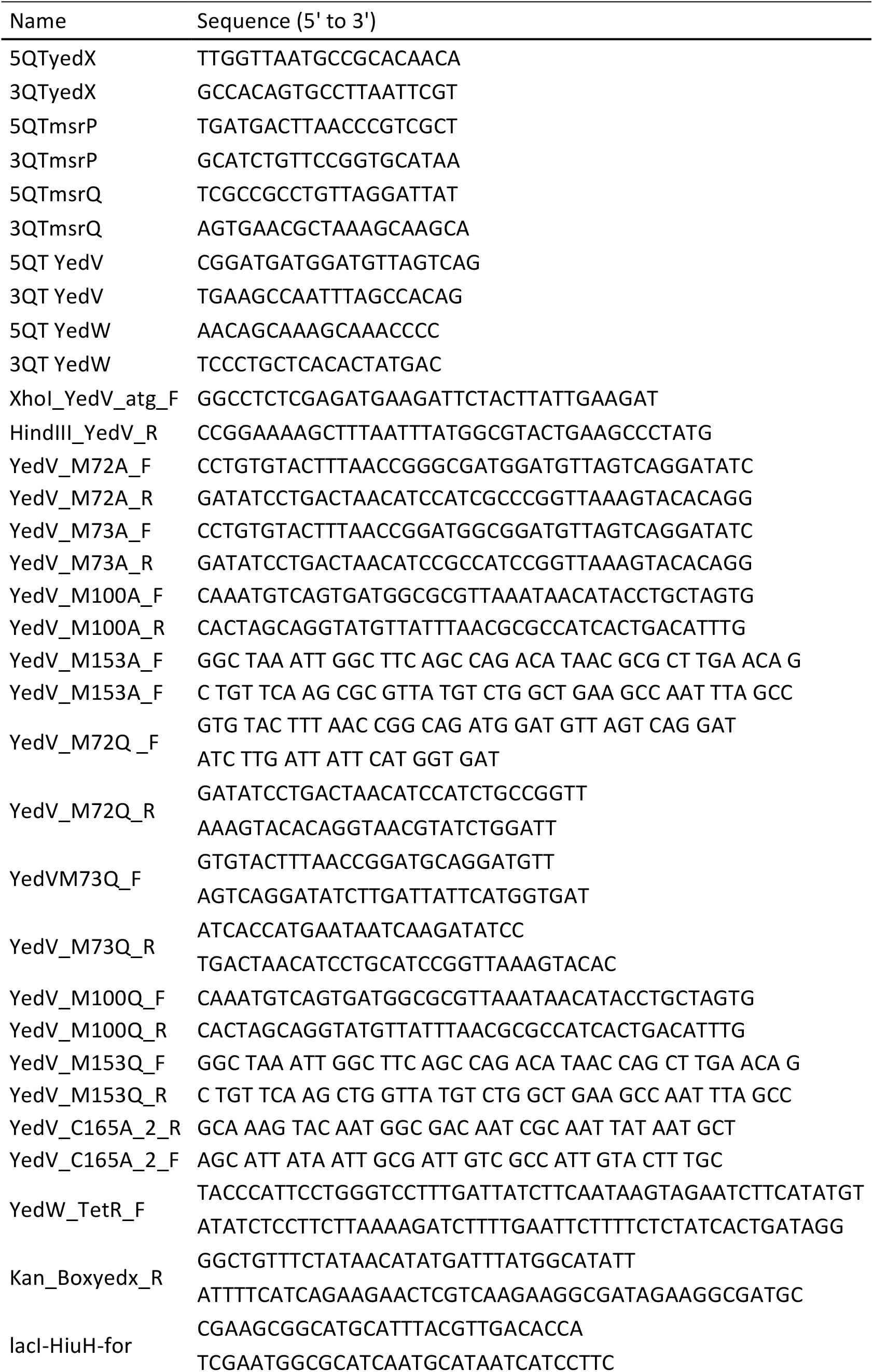
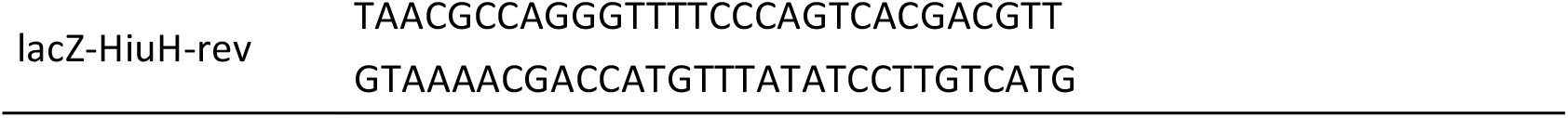
Primers used in this study. This table contains information regarding the primers used in this study, including their names and their sequence.

### Operonic organization analysis

*E. coli* cells (MG1655) were grown to OD_600_≈ 2 in the presence of 2 mM HOCl added at the beginning of the culture. Total bacterial RNA was extracted with Maxwell® 16 LEV miRNA Tissue Kit (Promega) according to the manufacturer’s instruction. Contaminating chromosomal DNA was digested using Turbo DNase (Invitrogene). RNA quality was assessed using Tape station system (Agilent) then quantified at 260 nm using NanoDrop 1000 (Thermo Fisher scientific). cDNA was synthesized with 1 µg of total RNA, 0.5 µg random primers (Promega) and GoScript™ reverse transcriptase (Promega). To analyze the *hiuH-msrP-msrQ* operon structure, two sets of primers, 5QTyedX-3QTmsrP and 5QTmsrP-3QTmsrQ, were used to amplify the possible transcripts. As a negative control, RNA samples were amplified using the same sets of primers in order to verify the absence of chromosomal contamination. As a positive control, chromosomal DNA was amplified using EconoTaq DNA polymerase (Promega). Results were visualized on a 1 % agarose gel prepared with TBE (Tris-Borate-EDTA) and stained with ethidium bromide.

### qRT-PCR analysis

As described previously, total RNA was extracted from wild type *E. coli* (MG1655) cells and *yedV* mutants (SH198) either cultured to OD_600_≈ 0.6 and subjected to 6 mM H_2_O_2_ for 30 min or cultured to OD_600_≈ 1 and stressed with 4 mM HOCl for 1 h. mRNA levels of *yedV, yedW, msrP* and *msrQ* were quantified by qRT-PCR analysis using the incorporation of EvaGreen (Bio-rad) in a CFX96 Real Time system (Bio-Rad) according to the manufacturer’s instruction. For these reactions, primers 5QT-3QT yedX, 5QT-3QT msrP, 5QT-3QT yedV, 5QT-3QT yedW were used. Amplification of 16S rRNA was used as a loading control. qRT-PCR technical triplicates for each condition were conducted. All biological repeats were selected and reported.

### HOCl induction assays

*E. coli* cells expressing *msrP-lacZ* or *hiuH-lacZ* (CH183, CH184 or CH186 carrying pYedV or pYedV derived plasmids) were grown in LB medium at 37 °C aerobically. At OD_600_≈ 1, the culture was divided into two 15 mL falcon tube containing each 5 mL of this culture. To one of those tubes, 4 mM HOCl (NaOCl, 10-15% AC Acros Organics) was added. Cultures were then incubated at 37 °C. To assess the activity of YedV mutants, β-galactosidase activity was measured after 1 h of incubation. To follow the kinetics of *msrP-lacZ* or *hiuH-lacZ* expression in response to HOCl, β-galactosidase activity was measured at different time points. For each protocol, the reaction mixture was prepared as follows: 200 µL of bacteria were added to 800 µL of β-galactosidase buffer. β-galactosidase was measured as previously described by Miller *et al*. ^31^.

### H_2_O_2_ induction assays

To test the induction of *msrP-lacZ* or *hiuH-lacZ* (CH183 and CH184) in response to H_2_O_2,_ two different protocols were tested: first, as described by Urano and his collaborators ^16^, *E. coli* cells were cultured in LB medium at 37 °C aerobically. At OD_600_≈ 0.6, cultures were split into two 50 mL falcon tubes each containing 5 mL of the culture. 6 mM of H_2_O_2_ was added to one of them. After 30 min of incubation at 37°C, β-galactosidase activity was measured. For the second protocol, *E. coli* cells were cultured in LB medium at 37°C aerobically. At OD_600_≈ 0.2, the culture was divided into 5 mL aliquot in 50mL falcon tubes. Different sub-lethal concentrations of H_2_O_2_ were added for each (500 µM, 1 mM, 2 mM and 3 mM of H_2_O_2_). 200 µL of culture were harvested at different time points and β-galactosidase assays were conducted. As a positive control for this protocol, strain BE1001 was used to follow the expression of *ahpC-lacZ* under the tested conditions.

### Bacteria survival assay in vitro

To test the effect of H_2_O_2_ and HOCl on cell viability, over night cultures of *E. coli* (MG1655) were diluted in LB medium to OD_600_≈ 0.4. Cells were then subjected to 6 mM H_2_O_2_ and 4 mM HOCl using the protocols described earlier. Cultures were serial-diluted in PBS. 5 µL of each dilution were spotted on LB agar plate. After 18h at 37 °C, CFU were counted.

### Chlorotaurine synthesis

Chlorotaurine (*N*-ChT) is produces by the reaction between HOCl and the amino acid taurine. Two identical volumes of 10 mM NaOCl and 10 mM taurine (Sigma) were mixed and incubated for 10 min at room temperature. Chlorotaurine concentration was then determined by measuring the solution absorbance at 252 nm (ε=429 M^-1^cm^-1^). Both HOCl and taurine solution were prepared in 0.1 M phosphate potassium buffer (pH 7.4).

### Chlorotaurine induction assays

To test the induction of *msrP-lacZ* by *N*-ChT, *E. coli* CH183 cells were grown in LB medium aerobically to OD_600_≈ 1. Then, 1 mM of *N*-ChT was added. 200 µL of culture were harvested at different time points and β-galactosidase activities were measured.

### NO induction assays

To test the induction of *msrP-lacZ by* NO, CH183 cells were grown in Hungate tubes in strictly anaerobic conditions with 10 mL LB medium at 37 °C. At OD_600_≈ 0.2, different concentrations of NO were applied (500 nM, 1 µM, 5 µM and 50 µM NO from a NO water saturated solution injected into the anaerobic cultures and followed by incubation at 37 °C. 200 µL of culture were harvested with a gastight Hamilton syringe at different time points and β-galactosidase activities were measured. Strain DV1301 expressing *hmpA-lacZ* fusion was used as a positive control.

### Superoxide radical induction assay

To test the induction of *msrP-lacZ* by superoxide radical, CH183 cells were grown in LB medium at 37 °C. At OD_600_≈ 0.2, the culture was split into 5 mL aliquots in 50 mL falcon tubes. Each culture was treated with different concentration of paraquat (300 µM, 500 µM and 1 mM paraquat) followed by incubation at 37 °C. 200 µL of culture were harvested at different time points and β-galactosidase activities were measured. BE1000 strain carrying *soxS-lacZ* was used as a positive control.

### Immunoblot analysis of MsrP expression

To assess MsrP induction by HOCl, overnight cultures of CH186 strain expressing different plasmidic alleles of *yedV* were diluted to OD_600_≈ 0.04 in LB medium. Cells were grown aerobically to OD_600_≈ 1. The culture was divided into two 15 mL falcon tube containing each 5 mL of the culture. To one of those tubes, 4 mM HOCl was added. Cultures were then incubated at 37 °C for 1 h. Samples were diluted in sample buffer (Laemmli: 2 % SDS, 10 % glycerol, 60 mM Tris-HCl, pH 7.4, 0.01 % bromophenol blue). The amount of protein loaded on the gel was standardized for each culture based on their A_600_ values. Samples were then heated for 10 min at 95 °C and loaded on SDS-PAGE gel for immunoblot analysis. Immunoblot analysis was preformed according to standard procedures: the primary antibody is a guinea pig anti-MsrP antibody (provided by Jean-François Collet Lab, De Duve Institute, Belgium). The secondary antibody is anti-guinea pig IgG conjugated to horseradish peroxidase (HRP) (Promega). ImageQuant LAS4000 camera (GE Healthcare Life Sciences) was used for image Chemiluminescence.

### Immunoblot analysis of YedV expression

To analyze the expression level of YedV mutants, overnight cultures of CH186 strain expressing different plasmidic alleles of yedV were diluted to OD_600_≈ 0.04 in LB medium. Cells were cultured for 4 hours at 37 °C. 1 mM of isopropyl β-D-1-thiogalactopyranoside (IPTG) was then added and cultures were incubated over-night at 37 °C. Preparation of samples was proceeded as described earlier. For immunoblot analysis, rabbit anti-YedV was used as a primary antibody (synthesized for Benjamin Ezraty Lab by Agro-bio) and an anti-rabit IgG conjugated to horseradish peroxidase (HRP) (Promega) was used as a secondary-antibody. Chemiluminescence signals were visualized using ImageQuant LAS4000 camera (GE Healthcare Life Sciences).

### Analysis of MsrP effect on hiuH expression

To analyze the effect of MsrP on the expression of *hiuH*, overnight cultures of SH100 and SH101 carrying or not plasmids pAG117 and pAG195 were diluted to OD_600_≈ 0.04 in M9 minimal media supplemented with 0.2 % Casaminoacids (Bio Basic Canada). Cultures were incubated at 37 °C. At OD_600_≈ 0.2, 135 µM of HOCl was added and β-galactosidase activity was measured at different time points as described earlier.

## ACKNOWLEDGEMENTS

We thank all the members of the Ezraty group for comments on the manuscript, advice and discussion throughout the work. We also thank the Py group (LCB-Marseille) and J.F. Collet (De Duve Institute-Brussels) for helpful suggestions and comments on this work. A special thank to the former Marseillaise Barras team (Team Barras 4 ever) and to Frederic Barras (now at Institut Pasteur) for lab space, support and discussion. This work was supported by grants from the Agence Nationale de la Recherche (ANR) (#ANR-16-CE11-0012-02 METOXIC), the Centre national de la recherche scientifique (CNRS) and Aix-Marseille Université (AMU).

---

**Supplementary Figure 1-.**
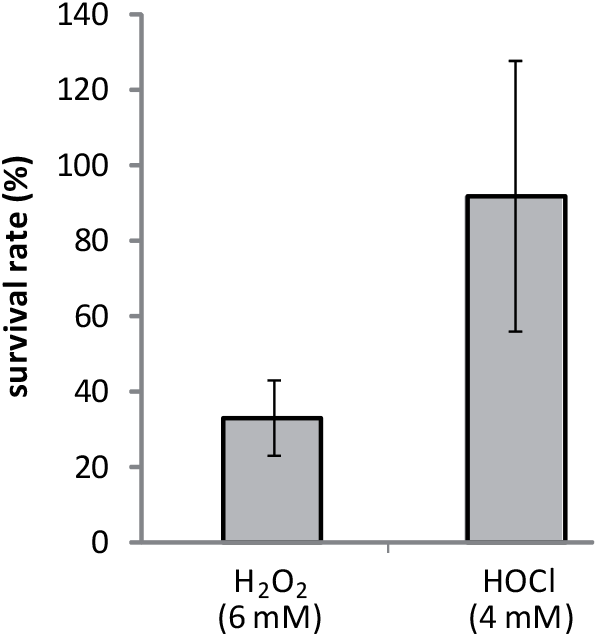
Effect of 6 mM H_2_O_2_ and 4 mM HOCl on the viability of WT *E. coli* cells. *E. coli* MG1655 (wild-type) strain were grown in LB medium. Each culture was subjected either to 6 mM H_2_O_2_ for 30 min^15^ or to 4 mM HOCl for 1h. Cultures were then serial diluted in PBS and spotted on LB agar plates. After 18 h of incubation, CFU were counted and compared to untreated cells. Values shown represent the ratio between CFU of treated cells/ CFU of untreated cells at dilution 10^−7^. The results are expressed as means ± SD of three independent experiments.

**Supplementary Figure 2-.**
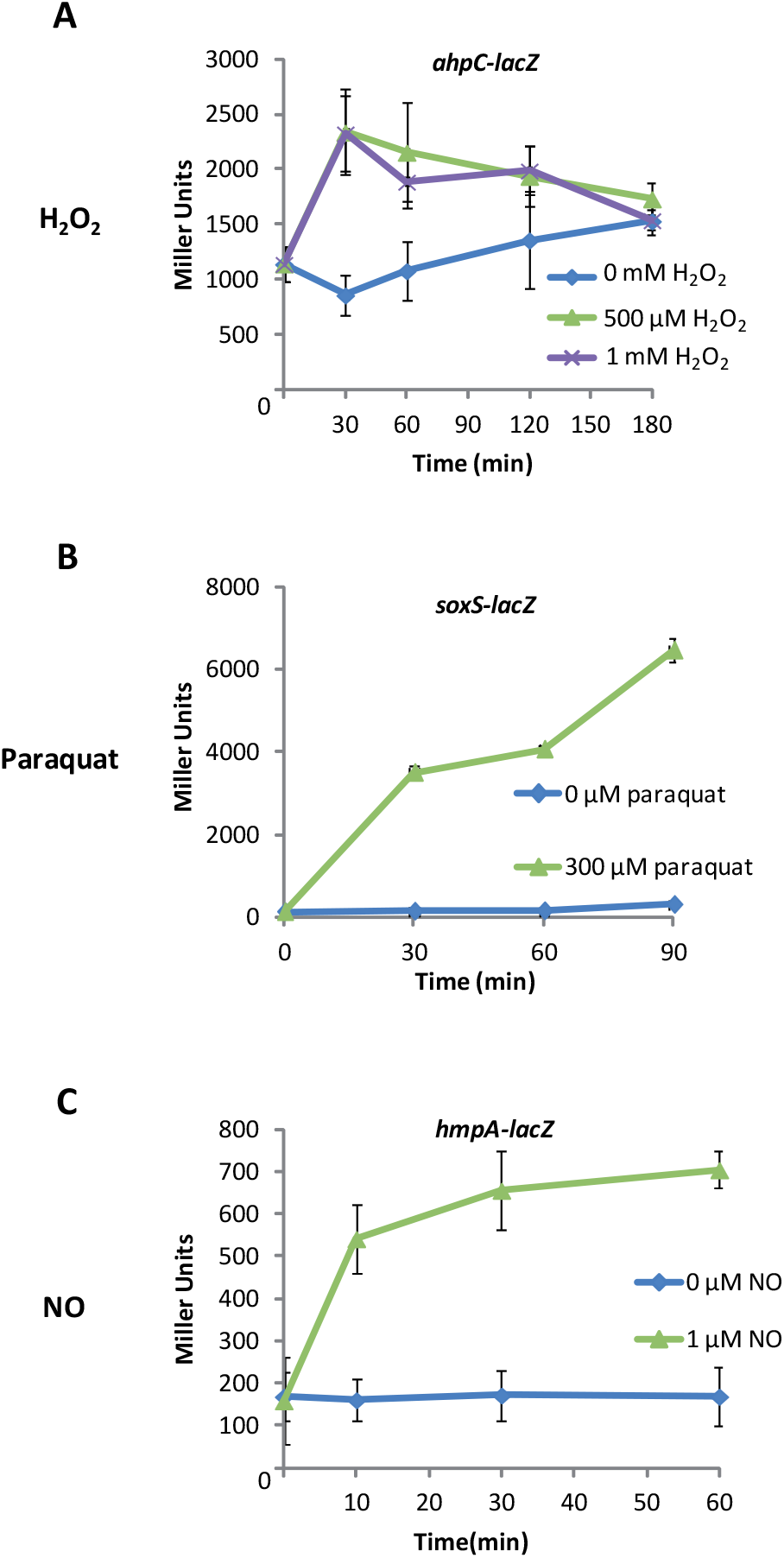
Strains used as a positive control in response to different oxidative stress. **A)** Effect of H_2_O_2_ on *ahpC-lacZ* expression. Strain BE1001 carrying *ahpC-lacZ* fusion, known to respond to H_2_O_2_, was subjected to sub-lethal concentrations of H_2_O_2_ (0.5 mM and 1 mM) and *ahpC-lacZ* expression was followed at different time points. Results are expressed as means ± SD (n=5) **B)** Effect of superoxide radical on *soxS-lacZ* expression. Paraquat was used to produce superoxide radical in the cells. The strain BE1000 carrying the fusion *soxS-lacZ*, known to respond to superoxide radical, was subjected to sub-lethal concentration of paraquat (300 µM) and the expression of *soxS-lacZ* was followed at different time points. Results are expressed as means ± SD (n=3) **C)** Effect of NO on *hmpA-lacZ* expression. The strain DV1301 carrying the fusion *hmpA-lacZ*, known to respond to NO, was subjected to sub-lethal concentration of NO (1 µM,). *hmpA-lacZ* expression was followed at different time points. Results are expressed as means ± SD (n=3)

## REFERENCES

1. Winterbourn, C. C., Hampton, M. B., Livesey, J. H. & Kettle, A. J. Modeling the Reactions of Superoxide and Myeloperoxidase in the Neutrophil Phagosome. J. Biol. Chem. 281, 39860–39869 (2006).

2. Hawkins, C. L. Hypochlorous acid-mediated modification of proteins and its consequences. Essays Biochem. 64, 75–86 (2019).

3. da Cruz Nizer, W. S., Inkovskiy, V. & Overhage, J. Surviving Reactive Chlorine Stress: Responses of Gram-Negative Bacteria to Hypochlorous Acid. Microorganisms 8, (2020).

4. Winterbourn, C. C. Comparative reactivities of various biological compounds with myeloperoxidase-hydrogen peroxide-chloride, and similarity of the oxidant to hypochlorite. Biochim. Biophys. Acta 840, 204–210 (1985).

5. Deborde, M. & von Gunten, U. Reactions of chlorine with inorganic and organic compounds during water treatment-Kinetics and mechanisms: a critical review. Water Res. 42, 13–51 (2008).

6. Winterbourn, C. C. Reconciling the chemistry and biology of reactive oxygen species. Nat. Chem. Biol. 4, 278–286 (2008).

7. Gray, M. J., Wholey, W.-Y. & Jakob, U. Bacterial responses to reactive chlorine species. Annu. Rev. Microbiol. 67, 141–160 (2013).

8. Drazic, A. et al. Methionine oxidation activates a transcription factor in response to oxidative stress. Proc. Natl. Acad. Sci. U. S. A. 110, 9493–9498 (2013).

9. Parker, B. W., Schwessinger, E. A., Jakob, U. & Gray, M. J. The RclR protein is a reactive chlorine-specific transcription factor in Escherichia coli. J. Biol. Chem. 288, 32574–32584 (2013).

10. Gray, M. J., Wholey, W.-Y., Parker, B. W., Kim, M. & Jakob, U. NemR is a bleach-sensing transcription factor. J. Biol. Chem. 288, 13789–13798 (2013).

11. Ezraty, B., Gennaris, A., Barras, F. & Collet, J.-F. Oxidative stress, protein damage and repair in bacteria. Nat. Rev. Microbiol. 15, 385–396 (2017).

12. Whitworth, D. E. & Cock, P. J. A. Evolution of prokaryotic two-component systems: insights from comparative genomics. Amino Acids 37, 459–466 (2009).

13. Gennaris, A. et al. Repairing oxidized proteins in the bacterial envelope using respiratory chain electrons. Nature 528, 409–412 (2015).

14. Lee, Y. et al. Transthyretin-related proteins function to facilitate the hydrolysis of 5-hydroxyisourate, the end product of the uricase reaction. FEBS Lett. 579, 4769–4774 (2005).

15. Urano, H., Umezawa, Y., Yamamoto, K., Ishihama, A. & Ogasawara, H. Cooperative regulation of the common target genes between H2O2-sensing YedVW and Cu^2+^-sensing CusSR in Escherichia coli. Microbiol. 161, 729–738 (2015).

16. Urano, H. et al. Cross-regulation between two common ancestral response regulators, HprR and CusR, in Escherichia coli. Microbiol. 163, 243–252 (2017).

17. Varatnitskaya, M., Degrossoli, A. & Leichert, L. I. Redox regulation in host-pathogen interactions: thiol switches and beyond. Biol. Chem. 402, 299–316 (2021).

18. Bigelow, D. J. & Squier, T. C. Thioredoxin-dependent redox regulation of cellular signaling and stress response through reversible oxidation of methionines. Mol. Biosyst. 7, 2101–2109 (2011).

19. Mizuno, T. Compilation of all genes encoding two-component phosphotransfer signal transducers in the genome of Escherichia coli. DNA Res. Int. J. Rapid Publ. Rep. Genes Genomes 4, 161–168 (1997).

20. Yamamoto, K. et al. Functional Characterization in Vitro of All Two-component Signal Transduction Systems from Escherichia coli. J. Biol. Chem. 280, 1448–1456 (2005).

21. Laub, M. T. The Role of Two-Component Signal Transduction Systems in Bacterial Stress Responses. in Bacterial Stress Responses 45–58 (John Wiley & Sons, Ltd, 2010). doi:10.1128/9781555816841.ch4.

22. Danchin, A. Coping with inevitable accidents in metabolism. Microb. Biotechnol. 10, 57–72 (2016).

23. Tung, Q. N., Busche, T., Van Loi, V., Kalinowski, J. & Antelmann, H. The redox-sensing MarR-type repressor HypS controls hypochlorite and antimicrobial resistance in Mycobacterium smegmatis. Free Radic. Biol. Med. 147, 252–261 (2020).

24. Loi, V. V. et al. Redox-Sensing Under Hypochlorite Stress and Infection Conditions by the Rrf2-Family Repressor HypR in Staphylococcus aureus. Antioxid. Redox Signal. 29, 615–636 (2018).

25. Buschiazzo, A. & Trajtenberg, F. Two-Component Sensing and Regulation: How Do Histidine Kinases Talk with Response Regulators at the Molecular Level? Annu. Rev. Microbiol. 73, 507–528 (2019).

26. Vogt, W. Oxidation of methionyl residues in proteins: tools, targets, and reversal. Free Radic. Biol. Med. 18, 93–105 (1995).

27. Baba, T. et al. Construction of Escherichia coli K-12 in-frame, single-gene knockout mutants: the Keio collection. Mol. Syst. Biol. 2, 2006.0008 (2006).

28. Bremer, E., Silhavy, T. J., Weisemann, J. M. & Weinstock, G. M. Lambda placMu: a transposable derivative of bacteriophage lambda for creating lacZ protein fusions in a single step. J. Bacteriol. 158, 1084–1093 (1984).

29. Datsenko, K. A. & Wanner, B. L. One-step inactivation of chromosomal genes in Escherichia coli K-12 using PCR products. Proc. Natl. Acad. Sci. U. S. A. 97, 6640–6645 (2000).

30. Mandin, P. & Gottesman, S. A genetic approach for finding small RNAs regulators of genes of interest identifies RybC as regulating the DpiA/DpiB two-component system. Mol. Microbiol. 72, 551–565 (2009).

31. Miller, J. H. A short course in bacterial genetics: a laboratory manual and handbook for Escherichia coli and related bacteria. (Cold Spring Harbor Laboratory Press, 1992).

32. Ezraty, B., Henry, C., Hérisse, M., Denamur, E. & Barras, F. Commercial Lysogeny Broth culture media and oxidative stress: a cautious tale. Free Radic. Biol. Med. 74, 245–251 (2014).

33. Vinella, D., Loiseau, L., Choudens, S. O. de, Fontecave, M. & Barras, F. In vivo [Fe-S] cluster acquisition by IscR and NsrR, two stress regulators in Escherichia coli. Mol. Microbiol. 87, 493–508 (2013).

34. Germain, E. et al. YtfK activates the stringent response by triggering the alarmone synthetase SpoT in Escherichia coli. Nat. Commun. 10, 5763 (2019).

